# Astrocytic synapse elimination controls ocular dominance plasticity

**DOI:** 10.1101/2022.09.23.509193

**Authors:** Joon-Hyuk Lee, Jinah Kim, Minjin Kim, Chi-Hong Shin, Won-Suk Chung

**Affiliations:** Department of Biological Sciences, and KAIST Stem Cell Center, Korea Advanced Institute of Science and Technology, Daejeon 34141, Republic of Korea

## Abstract

Ocular dominance plasticity (ODP) is a representative form of experience-dependent synaptic plasticity observed in the primary visual cortex (V1). However, the cellular mechanisms and physiological roles of synapse elimination in ODP are largely unknown. Here, we show that astrocytic phagocytosis of thalamo-cortical synapses in V1 is a critical mediator of ODP. We found that astrocytes, but not microglia, start to engulf thalamo-cortical synapses within 24 hours after monocular deprivation and that astrocytic synapse elimination is highly selective for synapses from the deprived eye, as revealed by AAV1-mediated trans-synaptic anterograde tracing of synapse phagocytosis reporters. Importantly, mice without the *Megf10* phagocytic receptor in astrocytes exhibit deficits in eliminating the synapses from the deprived eye, leading to the failure to reduce the number of thalamo-cortical synapses after monocular deprivation. Remarkably, *Megf10*-deficient animals show severe defects in monocular deprivation-induced cortical synapse remodeling and subsequent expansion of the thalamo-cortical circuitry from the nondeprived eye. Taken together, our data show that astrocytic synapse elimination through MEGF10 is one of the key components in ODP, revealing the physiological importance of astrocytic phagocytosis in experience-dependent synaptic plasticity.

## Main

Ocular dominance plasticity (ODP) is a classical model of developmental plasticity in which synaptic circuits are remodeled in response to alterations in sensory experiences by monocular deprivation (Wiesel and Hubel, 1965; Wiesel and Hubel, 1963a; Wiesel and Hubel, 1963; Chapman et al., 1986). To elucidate the cellular and molecular mechanisms underlying ODP, recent works have harnessed mice as a genetic model system (Gordon et al., 1996). In the mouse visual system, most of the projections from retinal ganglion cells (RGCs) are crossed at the optic chiasm and make synaptic connections with neurons in the contralateral lateral geniculate nucleus (LGN), while only a small portion of retinal inputs project to the ipsilateral LGN. This contra-versus ipsilateral connectivity between RGCs and the LGN makes a unique array of thalamo-cortical projections in the mouse primary visual cortex (V1), such that a large V1 area called the monocular zone (MZ) receives inputs only from the contralateral eye, whereas a relatively small V1 area called the binocular zone (BZ) receives input from both the contra- and ipsilateral eyes (Caviness et al., 1975). Upon visual deprivation of one eye, the brain starts to cope with this sensory experience by reorganizing the strength and connectivity of thalamo-cortical inputs in V1 (Gordon and Stryker, 1996), resulting in the shrinkage of the MZ and compensatory expansion of the BZ area (Tagawa et al., 2005). Interestingly, these changes are mostly detectable at the critical time window between postnatal Day 21 (P21) and P35 (Tagawa et al., 2005), while the adult brain shows limited ODP (Wiesel and Hubel, 1963; Tagawa et al., 2005). Decades of work have led to the identification of critical factors in mediating physiological synapse remodeling, such as synaptic depression and potentiation during ODP (Datwani et al., 2009; Syken et al., 2006; Katz et al., 1996); however, the cellular/molecular mechanisms of structural synapse elimination and its impact on the subsequent synapse formation process are largely unknown (Sawtell et al., 2003; Sun et al., 2019; Zhou et al., 2017; Oray et al., 2004; Van Versendaal et al., 2012).

Recent studies have revealed the nonautonomous mechanisms of synapse elimination by glial cells. Among them, microglia, CNS-resident immune cells, have been shown to mediate developmental synapse pruning (Wu et al., 2015; Stevens et al., 2007; Schafer et al., 2012) as well as disease-related synapse loss (Vasek et al., 2016; Lui et al., 2016; Hong et al., 2016) by phagocytosing synapses through the classical complement cascade. In addition, CX3CR1, a chemokine fractalkine receptor in microglia (Nishiyori et al., 1998), plays important roles in microglia-dependent circuit remodeling in the developing hippocampus (Paolicelli et al., 2011) as well as in the barrel cortex after whisker lesioning (Gunner et al., 2019). However, it has been shown that, surprisingly, blocking either the complement (Welsh et al., 2020) or fractalkine pathways (Schecter et al., 2017) in microglia does not interfere with monocular deprivation-induced synaptic reorganization in V1, suggesting the possibility of another mediator in this process.

Recent studies have identified astrocytes as critical components in synapse elimination during developmental as well as adult stages in healthy and diseased brains. Astrocytes are known to mediate the pruning of RGC inputs in the LGN for developmental eye-specific retinogeniculate segregation through the MEGF10 and MERTK phagocytic pathways (Chung et al., 2013). In adult brains, it has also been shown that constant clearance of hippocampal synapses by MEGF10-dependent phagocytosis of astrocytes is crucial for maintaining functional and structural homeostasis of the hippocampus, suggesting its physiological importance in the post developmental synapse remodeling (Lee et al., 2021). Interestingly, a previous classical work showed that transplanting primary astrocytes from young kitten to the V1 of adult cat was able to reinduce the ODP (Muller and Best, 1989), implicating potential roles of astrocytes in modulating V1 synaptic plasticity. Indeed, several studies have shown that astrocytes can respond to monocular deprivation by changing their shape (Hawrylak and Greenough, 1995) and releasing synaptogenic molecules, such as Hevin, to re-enforce thalamo-cortical synapses (Singh et al., 2016). However, how astrocytes contribute to the structural elimination of synapses during ODP is completely unknown.

Here, we show that astrocytes, but not microglia, start to engulf thalamo-cortical synapses in layer 4 of V1 within 24 hours after monocular deprivation. This sudden increase in synapse elimination by astrocytes is highly specific to synapses originating from the sensory-deprived eye, as revealed by AAV1-mediated transsynaptic anterograde tracing. Furthermore, we found that astrocytic synapse elimination is regulated by input-specific neural competition in V1 but not by general dampening of neural activity. Importantly, we show that astrocytes actively respond to monocular deprivation by increasing the mRNA expression of a *Megf10* phagocytic receptor. Mice without *Megf10* specifically in astrocytes fail to eliminate thalamo-cortical synapses in V1 after monocular deprivation, leading to failure to reduce the number of thalamo-cortical synapses. Remarkably, *Megf10*-deficient animals show severe defects in monocular deprivation-induced cortical synapse remodeling and expansion of the thalamo-cortical circuitry from the nondeprived eye. Taken together, our data show that astrocytic synapse elimination through MEGF10 is one of the key components in ODP, revealing the physiological importance of astrocytic phagocytosis in controlling experience-dependent structural synaptic plasticity.

## Results

### Basal levels of synapse engulfment by glial cells in the primary visual cortex

To investigate the potential contribution of glia-mediated synapse phagocytosis in synapse remodeling after monocular deprivation, we labeled the LGN-derived thalamo-cortical synapses with synapse phagocytosis reporters that we have previously developed (Lee et al., 2021). Briefly, these reporters utilize different pKa values of mCherry and eGFP, such that mCherry fluorescence is preserved and the EGFP signal is lost in low-pH lysosomes.

By injecting AAV9-packaged human Synapsin promoter (hSyn)-driven synaptophysin-mCherry-eGFP (AAV9::hSyn-syn1-mCherry-eGFP or AAV9::ExPre) into the LGN (Figure 1A), we first confirmed that LGN-derived thalamo-cortical synapses in layer 4 of V1 were labeled with mCherry-eGFP (Figures 1B-1C). When we stained AAV9-injected WT V1 with presynaptic markers, including VGluT2 (for thalamo-cortical presynapses), VGluT1 (for cortico-cortical presynapses) and VGAT (for inhibitory presynapses), we found that LGN-derived mCherry-eGFP expression was exclusively confined to VGluT2-positive thalamo-cortical synapses in layer 4 of V1, validating the specific targeting of presynaptic mCherry-eGFP to cortical LGN-derived synapses (Figure 1D-1F).

**Figure 1.**
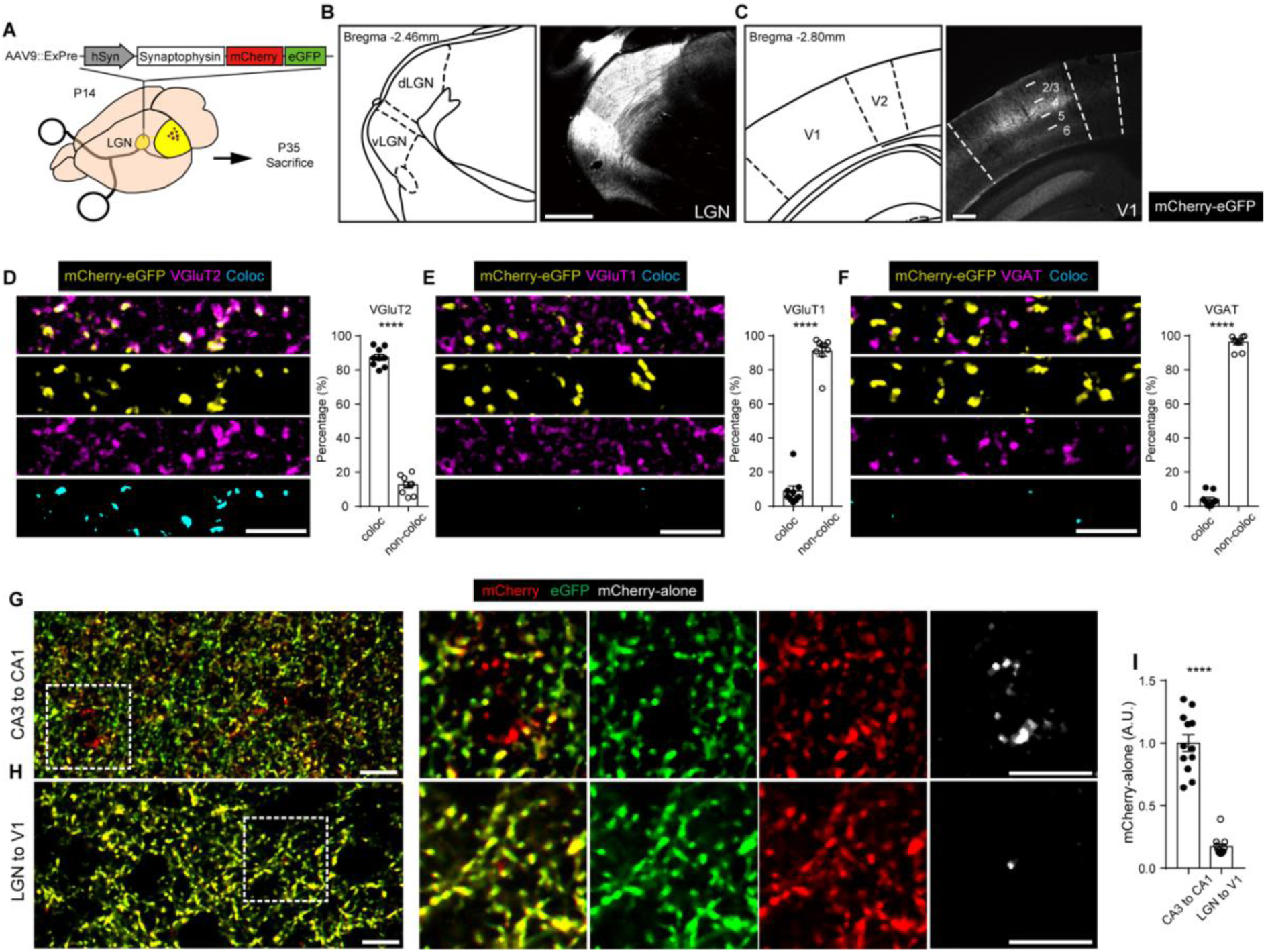
Basal levels of synapse engulfment by glial cells in the primary visual cortex. **(A)** Schematic illustration of the experimental design. All mice at P14 were injected with AAV9::ExPre into the LGN. After 3 weeks (at P35), all mice were sacrificed. **(B-C)** Representative images of mCherry-eGFP-expressing (white pseudocolored) LGN (B) and V1 (C) of AAV9::ExPre-injected mice. (Left) Anatomical location of the LGN (B, Bregma −2.46 mm) and V1 (C, Bregma −2.80 mm) based on the brain atlas. (Right) The numbers in the right panel of (C) indicate the layers of V1. Scale bar = 500 µm (B) and 250 µm (C). **(D-F)** (Left) Representative single plane images of mCherry-eGFP-labeled synapses (yellow) in V1 with complementary synaptic staining (D: VGluT2, E: VGluT1, F: VGAT, magenta). Colocalized synaptic puncta are shown in cyan. Scale bars = 5 µm. (Right) Percentage area of colocalized (coloc) and non colocalized (noncoloc) mCherry-eGFP-labeled synapses with complementary synaptic staining, which are further normalized by total mCherry-eGFP-labeled synapses. n=9 per group from 3 mice. Mean ± s.e.m. For all comparisons, ****P<0.0001, Mann‒Whitney t test. **(G-H)** Representative single plane images of mCherry-eGFP expression in CA1 (G) at P35 after CA3 injection of AAV9::ExPre at P14 and V1 (H) at P35 after LGN injection of AAV9::ExPre at P14. (Right) Magnified views of white dotted squares in the left panels. Scale bars = 10 µm. **(I)** Quantified areas of mCherry-only puncta in CA1 and V1 normalized to total mCherry-eGFP-labeled synapses. n=12 per group from 3 mice. Mean ± s.e.m. ****P<0.0001, Mann‒Whitney t test.

Previously, it has been suggested that after developmental synapse remodeling, neocortical synapses exhibit a long half-life compared to synapses in the hippocampal CA1 regions, where most synapses are turned over within several weeks (Attardo et al., 2015). Therefore, we examined the basal levels of glia-mediated synapse engulfment in V1 in comparison with those in the hippocampal CA1 by injecting AAV9::ExPre into the hippocampal CA3 or LGN of postnatal day (P) 14 mice. When we monitored CA1 and V1 areas at P35 to evaluate the level of mCherry-only puncta and putative synapse engulfment by glial cells, we found that compared to the numerous mCherry-only puncta in the hippocampal CA1, the level of mCherry-only puncta in V1 was significantly lower (Figure 1G-1I). These data indicate that thalamo-cortical synapses in V1 at juvenile stages exhibit very little glia-mediated synapse elimination, presumably reflecting a slow rate of synapse turnover in the cortical regions.

### Monocular deprivation induces glial phagocytosis of thalamo-cortical synapses

Although basal phagocytic activity of glial cells within V1 is extremely low, we hypothesized that depriving sensory stimulation in one eye (monocular deprivation) could induce functional changes in neuro-glial interactions, enabling glial cells to initiate the removal of synapses from the deprived eye. To address our hypothesis, we injected AAV9::ExPre into the left LGN of P14 mice and closed the right eye by suturing the eyelid at P32 (Figure 2E) so that monocular deprivation-mediated ODP could be initiated within the critical time window. Surprisingly, we observed a dramatic increase in the level of mCherry-only puncta within layer 4 of contralateral V1 at 1 and 2 days after monocular deprivation, which started to level off at 3 days (Figure 2F-G). These data indicate that glial cells in V1 start to engulf thalamo-cortical synapses soon after monocular deprivation is initiated, and this robust phagocytosis of synapses persists for at least 3 days during ODP-mediated synapse remodeling.

**Figure 2.**
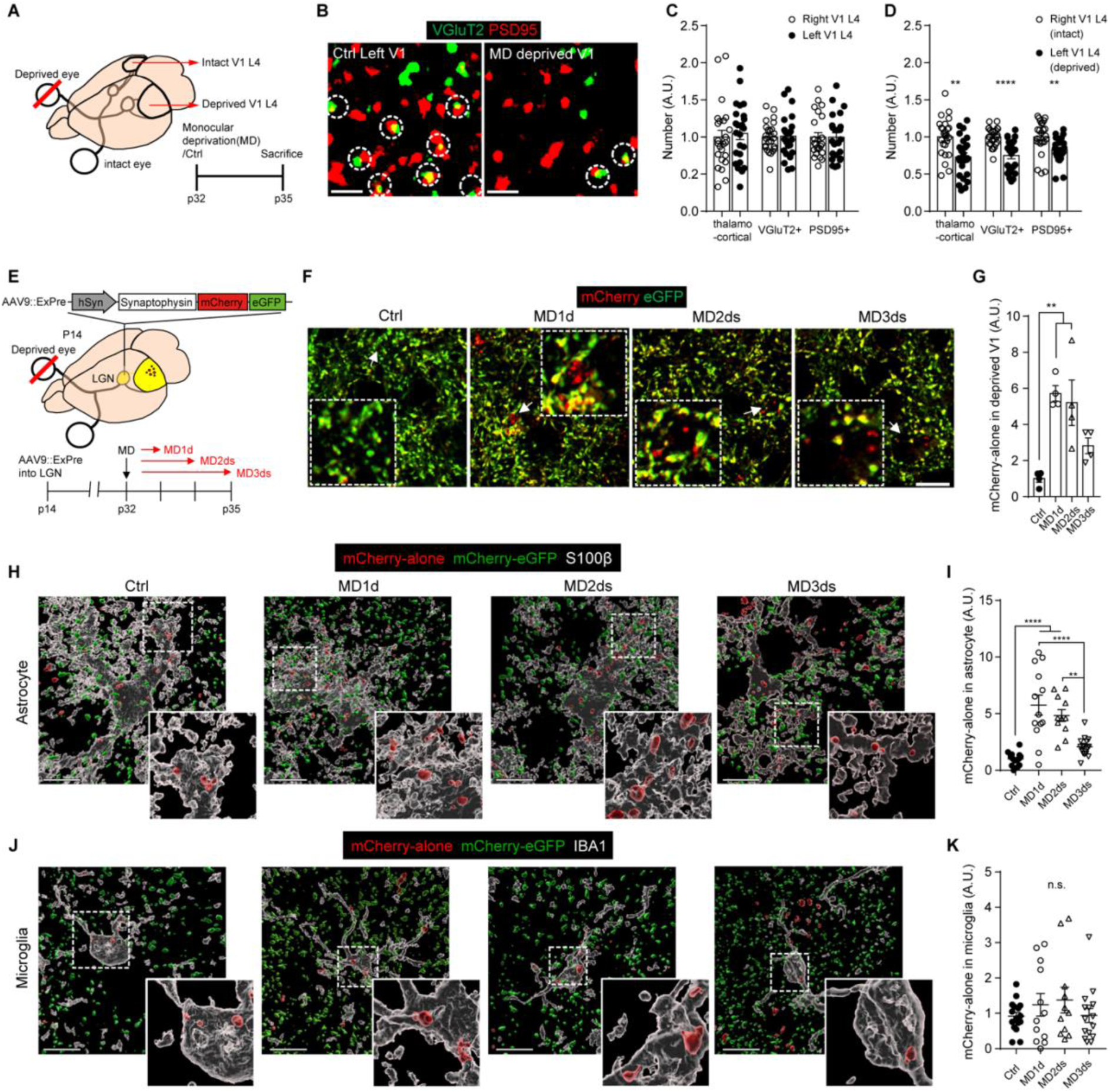
Monocular deprivation induces glial phagocytosis of thalamo-cortical synapses. **(A)** Schematic illustration of the experimental design and schedule. The right eyes of P32 mice were left intact (Ctrl) or deprived (MD), and the mice were sacrificed at P35. Left V1 (deprived) and right V1 (intact) were compared. **(B)** Representative single-plane images of thalamo-cortical excitatory synapses (VGluT2, green and PSD95, red) in V1 of the left hemisphere of the Ctrl (Ctrl Left V1) and MD (MD deprived V1) groups. White dotted circles indicate colocalized synapses. Scale bars = 1 µm. **(C-D)** Quantified number of thalamo-cortical, VGluT2+ and PSD95+ synapses in V1 of the Ctrl (C) and MD (D) groups. n=24 per group from 4 mice. Mean ± s.e.m. (C) For all groups, P>0.05, n.s., not significant. (D) thalamo-cortical, **P=0.0011; VGluT2+, ****P<0.0001; PSD95+, **P=0.0012; Mann‒Whitney t test. **(E)** Schematic illustration of the experimental design and schedule. Mice at P14 were injected with AAV9::ExPre into the LGN. The right eyes of P32 mice were kept intact (Ctrl) or deprived (MD), and the mice were sacrificed 1, 2 or 3 days after monocular deprivation (MD1d, MD2ds and MD3ds, respectively). **(F)** Representative single plane images of mCherry-eGFP expression in V1 of the Ctrl, MD1d, MD2ds and MD3ds groups. Inset white dotted square regions show magnified views of areas indicated by arrowheads. Scale bars = 10 µm. **(G)** Quantified areas of mCherry-only puncta in V1 normalized to total mCherry-eGFP-labeled synapses. n=4 mice per group; mean ± s.e.m.; (Ctrl vs. MD1d), **P=0.0024; (Ctrl vs. MD2ds), **P=0.0058. Data were compared using one-way ANOVA followed by multiple comparisons. **(H, J)** Representative 3D-rendered images showing ExPre-derived mCherry-only puncta and mCherry-eGFP puncta associated with astrocytes (H, S100b) and microglia (J, IBA1) in the Ctrl, MD1d, MD2ds and MD3ds groups. Lower-right panels are magnified views of white dotted square regions. Scale bars = 10 µm. **(I, K)** Quantified areas of mCherry-only puncta in astrocytes (I) and microglia (K) in V1 normalized to total mCherry-eGFP-labeled synapses. n=11 to 16 from 4 mice per group; mean ± s.e.m. For (I), (Ctrl vs. MD1d, MD2ds), ****P<0.0001; (MD1d vs. MD3ds), ****P<0.0001; (MD2ds vs. MD3ds), **P=0.0010. For all comparisons in (K), P>0.05, n.s., not significant. Data were compared using one-way ANOVA followed by multiple comparisons.

Next, to determine whether the increased glial phagocytosis is correlated with the reduced synapse number, we immunostained V1 after 3 days of monocular deprivation for VGluT2 and PSD95 and then compared the number of excitatory thalamo-cortical synapses between the contralateral and ipsilateral V1 (Figure 2A). As previously shown, the number of VGluT2-positive presynapses and PSD95-positive postsynapses as well as their synaptic contacts (VGluT2/PSD95 double-positive) were significantly reduced in layer 4 of the contralateral V1 compared to layer 4 of the ipsilateral V1 after 3 days of monocular deprivation (Figure 2B-2C). Interestingly, when we measured and compared the numbers of VGluT1-positive excitatory presynapses as well as VGAT/Gephyrin-positive inhibitory synapses in layer 4 of V1, their numbers were not changed between the contralateral and ipsilateral V1 in the mice after 3 days of monocular deprivation (Figure S1A-S1B). These data indicate that monocular deprivation primarily affects the number of excitatory thalamo-cortical synapses in layer 4 of V1.

### A selective increase in astrocytic phagocytosis of thalamo-cortical synapses after monocular deprivation

Since both astrocytes and microglia possess phagocytic capacity and participate in synapse pruning to sculpt and refine the visual circuit in the developing retinogeniculate system (Stevens et al., 2007; Schafer et al., 2012; Chung et al., 2013), we investigated which glial cells primarily eliminate thalamo-cortical synapses in V1 after monocular deprivation. To address this question, we induced monocular deprivation in mice injected with AAV9::ExPre into the LGN and immunostained astrocytes and microglia in V1 after 1∼3 days with antibodies for S100b (Figure S2) or IBA1 (microglia). To our surprise, we found that a substantial number of mCherry-only puncta was localized in S100b-positive astrocytes within layer 4 of V1 at 1 and 2 days after monocular deprivation, which started to be reduced at 3 days (Figure 2H-2I), similar to our ExPre data without glial staining (Figure 2G). Interestingly, although IBA1-positive microglia do contain the similar amount of baseline mCherry-only puncta compared to astrocytes (Figure S3), it was not increased by monocular deprivation (Figure 2J-2K), suggesting that microglia do not participate in eliminating thalamo-cortical synapses in response to monocular deprivation.

To determine whether excitatory postsynapses in layer 4 of V1 are also eliminated by glial phagocytosis after monocular deprivation, we injected the excitatory postsynapse-specific mCherry-eGFP reporter (AAV9::hSyn-driven PSD95Δ1,2-mCherry-eGFP, or AAV9::ExPost2) into V1 at P14, closed the right eyelid at P32 to induce monocular deprivation and sacrificed the mice after 2 days (Figure S4A-S4D). Consistent with our data from the ExPre reporter, we found that the number of ExPost2-derived mCherry-only puncta inside V1 astrocytes was significantly increased after 2 days of monocular deprivation compared to that in the control (Figure S4E). However, microglia did not show significant changes in the number of ExPost2- derived mCherry-only puncta after 2 days of monocular deprivation compared to the control level (Figure S4F).

Together, these data suggest that astrocytes, but not microglia, actively participate in monocular deprivation-dependent elimination of thalamo-cortical synapses by phagocytosing excitatory pre- and postsynapse compartments. The absence of alterations in microglial phagocytosis after monocular deprivation is also consistent with a previous report that microglial complement and fractalkine pathways do not play major roles in ODP-related synapse phagocytosis (Welsh et al., 2020; Schecter et al., 2017).

### Astrocytes preferentially eliminate the deprived thalamo-cortical synapses

V1 receives thalamo-cortical inputs from the LGN connected with both contralateral and ipsilateral eyes. Instead of forming an ocular dominance column that can be found in the V1 of higher mammals, the mouse V1 is anatomically segregated into the MZ, where V1 neurons receive inputs only from the contralateral eye, or the BZ, where V1 neurons receive inputs from both the contralateral and ipsilateral eyes (Caviness, 1975). To distinguish contralateral versus ipsilateral eye-derived V1 thalamo-cortical synapses in the same V1 hemisphere, we employed AAV1-mediated transsynaptic anterograde tracing, which takes advantage of the ability of AAV1 to be anterogradely spread to downstream neurons (Zingg et al., 2017). To verify the AAV1-mediated tracing system in LGN to V1 circuits, we injected AAV1-hSyn-Cre into the contra- or ipsilateral eye in conjunction with AAV9::EF1alpha double-floxed inverse open reading frame (DIO)-eYFP injection into the left LGN of P14 mice. Three days before sacrifice, cholera toxin subunit B-conjugated Alexa 594 fluorophore (CTB594) was injected into the contralateral eye to label the LGN domain that receives the contralateral eye-derived synapses (Figure S5A, S5C). Notably, we found that our AAV1-mediated transsynaptic anterograde tracing method enabled us to label selective subpopulations of LGN neurons that receive either contralateral or ipsilateral eye-derived synaptic connections (Figure S5B, S5D).

By taking advantage of this system, we succeeded in expressing the ExPre reporter in thalamo-cortical synapses in an eye-specific manner by injecting AAV1-hSyn-Cre into the contralateral or ipsilateral eye and AAV9::hSyn-DIO-ExPre into the left LGN (Figure 3A-3C). Sparsely labeled thalamo-cortical presynapses with mCherry-eGFP were both located within layer 4 of the left V1 (Figure 3B-C). Importantly, when we measured the number of mCherry-only puncta inside astrocytes after 2 and 3 days of monocular deprivation, we found that the number of mCherry-only puncta derived from contralateral (deprived) eye-specific synapses was significantly increased in V1 astrocytes (Figure 3D). However, the number of mCherry-only puncta derived from ipsilateral (nondeprived) eye-specific synapses showed similar levels in V1 astrocytes after monocular deprivation (Figure 3E). These data strongly indicate that astrocytes can recognize and preferentially eliminate synapses originating from the sensory-deprived eye while preserving synapses from the normal eye after monocular deprivation.

**Figure 3.**
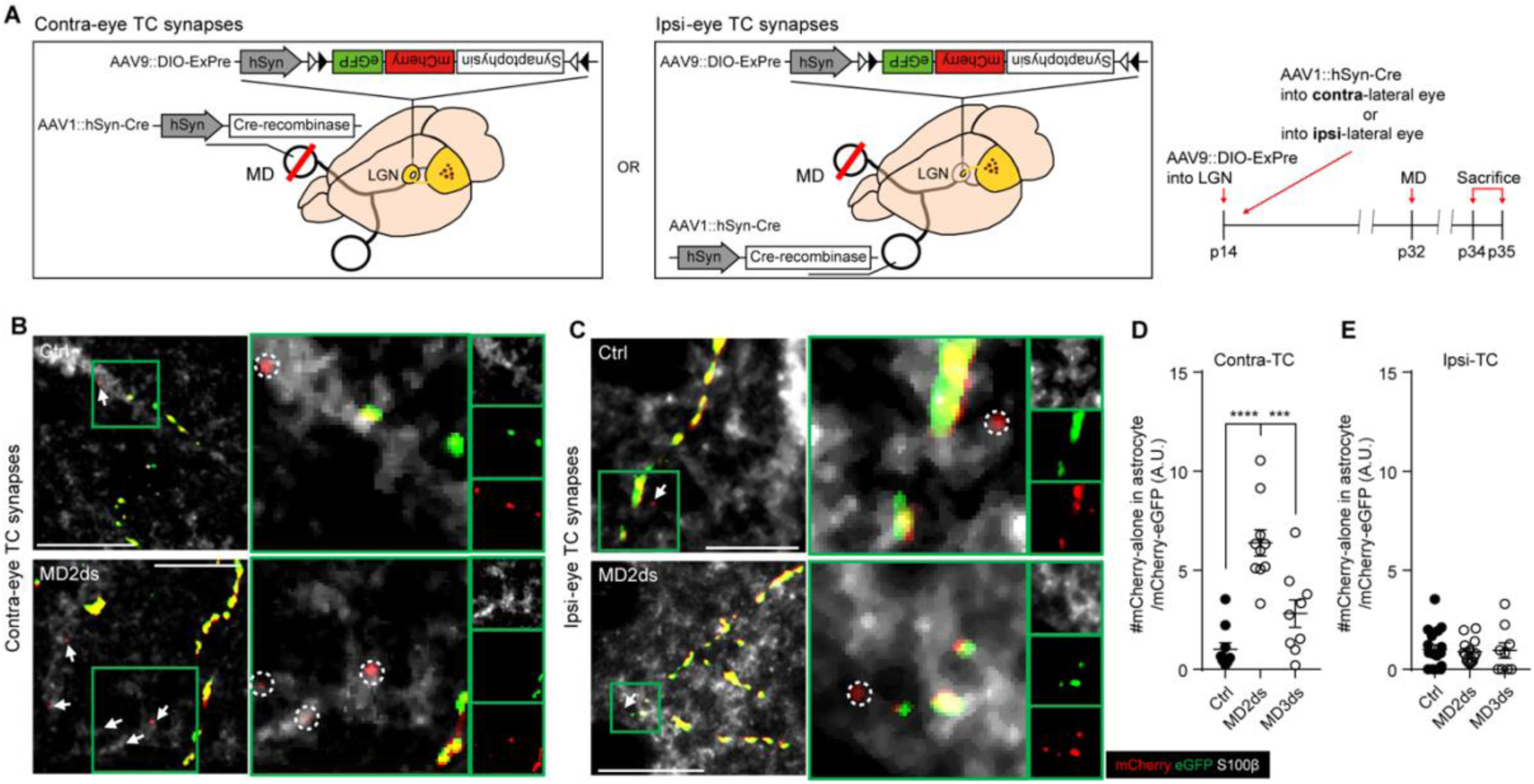
Astrocytes preferentially eliminate deprived thalamo-cortical synapses. **(A)** Schematic illustration of the experimental design and schedule. All mice at P14 were injected with AAV9::DIO-ExPre into the LGN and AAV1::hSyn-Cre into the right (contralateral) or left (ipsilateral) eye. The right eyes of P32 mice were kept intact (Ctrl) or deprived (MD), and the mice were sacrificed at P34 or P35 (MD2ds or MD3ds, respectively). **(B-C)** Representative z-stacked images showing ExPre-derived mCherry and eGFP puncta associated with astrocytes (S100b) in intact (Ctrl) or deprived (MD2ds) V1 of the contralateral eye (B) or ipsilateral eye (C) thalamo-cortical synapse group. Right panels highlight the green square-indicated area on the left panels. White arrows and dotted circles indicate mCherry-only puncta inside astrocytes. Scale bars = 10 µm. **(D-E)** Quantified number of mCherry-only puncta in astrocytes in V1 of the contralateral eye (D) or ipsilateral eye (E) thalamo-cortical synapse group normalized by total mCherry-eGFP-labeled synapses. n=10-17 from 3-4 mice per group. Mean ± s.e.m. For (D), (Ctrl vs. MD2ds), ****P<0.0001; (MD2ds vs. MD3ds), ***P=0.0006. For all comparisons in (E), P>0.05, n.s., not significant. Data were compared using one-way ANOVA followed by multiple comparisons.

### Neural activity-dependent competition drives thalamo-cortical synapse elimination by astrocytes

Previous studies have shown that astrocytic phagocytosis can be regulated by neural competition in the developing retinogeniculate system (Chung et al., 2013). To determine whether monocular deprivation induces astrocytic synapse elimination by potentiating neural competition between synapses derived from intact versus deprived eyes, we compared the extent of astrocyte-mediated elimination of thalamo-cortical synapses after monocular or binocular deprivation. Contrary to monocular deprivation, binocular deprivation has been known to dampen general neural activity in V1, diminishing synaptic competition and ODP (Hubel and Wiesel, 1965). After inducing 2 days of monocular or binocular deprivation in the mice injected with AAV9::Expre into the LGN at P14 (Figure 4A), we found that, in sharp contrast to the results from monocular deprivation, binocular deprivation failed to increase astrocytic mCherry-only puncta derived from thalamo-cortical synapses (Figure 4B-4C). These data indicate that astrocyte-mediated phagocytosis of synapses is initiated not by dampening neural activity, but by the increase in synaptic competition. Interestingly, we also noticed that the number of astrocytic mCherry-only puncta found in the MZ and BZ was not significantly different upon monocular deprivation, suggesting that there was an overall increase in active synapse remodeling by astrocytic phagocytosis in the MZ and BZ V1 regions during ODP (Figure 4D).

**Figure 4.**
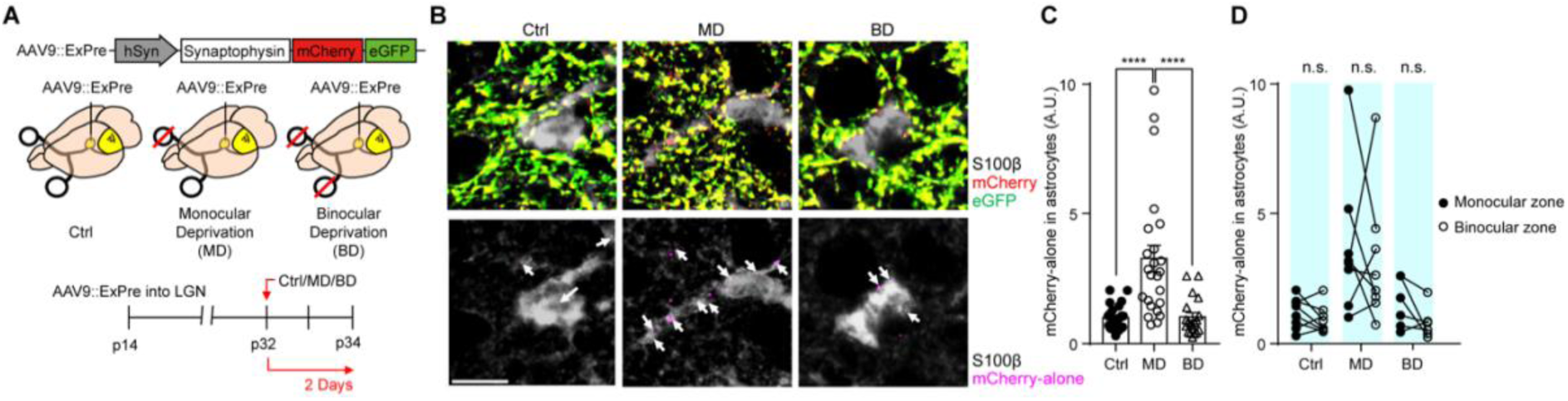
Neural activity-dependent competition drives thalamo-cortical synapse elimination by astrocytes. **(A)** Schematic illustration of the experimental design and schedule. All mice at P14 were injected with AAV9::ExPre into the LGN. For the Ctrl group, both eyes remained intact. For the MD group, the right eyes of P32 mice were removed. For the BD group, both eyes of P32 mice were deprived. All mice were sacrificed at P34. **(B)** Representative z-stacked images showing ExPre-derived mCherry and eGFP puncta associated with astrocytes (S100b) in V1 of the Ctrl, MD and BD groups. White arrows in the lower panels indicate mCherry-only puncta (magenta) inside astrocytes. Scale bars = 10 µm. **(C)** Quantified areas of mCherry-only inside astrocytes in V1 normalized to total mCherry-eGFP-labeled synapses. n=19-24 from 4 mice per group. Mean ± s.e.m.; (Ctrl vs. MD), ****P<0.0001; (MD vs. BD), ****P<0.0001. Data were compared using one-way ANOVA followed by multiple comparisons. **(D)** Comparisons of quantified mCherry-only areas inside astrocytes between the monocular zone and binocular zone within each group (Ctrl, MD and BD). n=19-24 from 4 mice per group. Mean ± s.e.m. For all comparisons, P>0.05, n.s., not significant. Mann‒Whitney t test.

### Monocular deprivation induced astrocytic gene expression changes

Next, to investigate the potential gene expression changes in V1 astrocytes after monocular deprivation, we performed bulk RNA-seq experiments using magnetic-activated cell sorting (MACS)-purified astrocytes (Figure S6A, B) from the P34 V1 after inducing monocular deprivation for 2 days (Figure 5A-B). We found that many astrocytic genes were differentially affected by monocular deprivation, including *C1qa, b, c* and *Egr1* (Figure 5C), indicating that astrocytes actively respond to changes in V1 synapse remodeling induced by monocular deprivation. For example, V1 astrocytes after monocular deprivation showed an increase in genes related to microtubule bundle formation, axoneme assembly, and the PI3K-Akt pathway (Figure 5D) and a decrease in genes related to chromatin remodeling, the complement cascade, and the regulation of immune system pathways (Figure 5E). To directly compare the changes in phagocytic gene expression in astrocytes, we analyzed the z scored Reads Per Kilobase of transcript, per Million mapped reads (RPKM) value of phagocytic receptors, such as *Megf10*, *Mertk* and *Axl*, which have been shown to be expressed by astrocytes. Interestingly, we found that the mRNA expression of *Megf10* was significantly increased after monocular deprivation (Figure 5F), whereas expression of *Mertk* and *Axl* did not show this increase (Figure 5G-H). In line with these data, we found that the immunoreactivity of anti-MEGF10 in astrocytes was significantly increased (Figure 5I-5J), while that of anti-MERTK or anti-AXL was not changed after monocular deprivation (Figure 5K-5N). We did not observe changes in the mRNA levels of genes related to astrocytic synaptogenesis (*Thbs1*, *sparcl* and *gpc4*) in our experimental setting (Figure S6C-6E). Since our RNA-seq data strongly suggest that MEGF10 can be a mediator of thalamo-cortical synapse elimination by astrocytes, we decided to interrogate the role of astrocytic MEGF10 in selectively eliminating and remodeling thalamo-cortical synapses during ODP.

**Figure 5.**
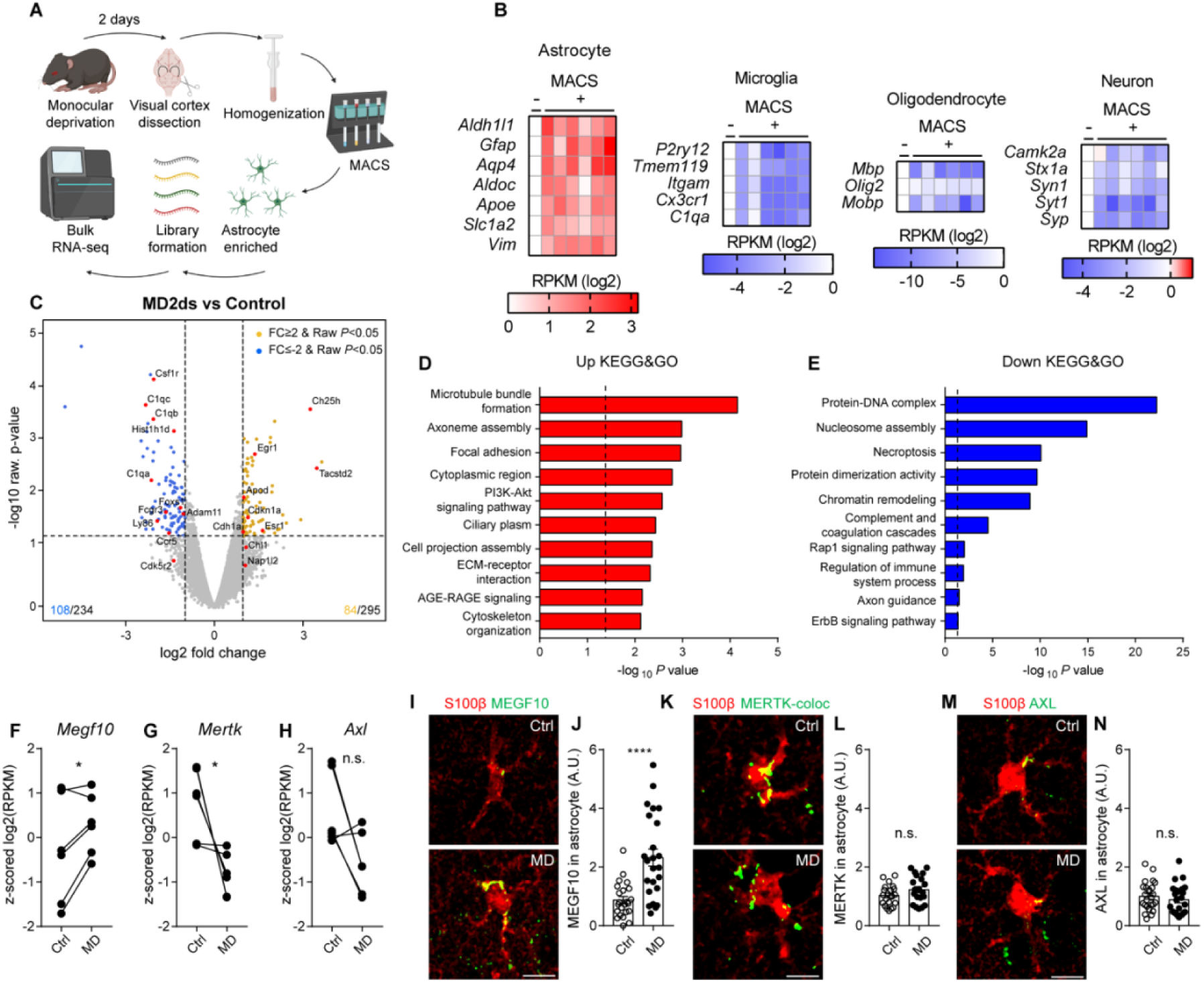
Monocular deprivation induces gene expression changes in astrocytes. **(A)** Schematic illustration of the experimental design. The right eyes of P32 mice remained intact (Ctrl) or were deprived for 2 days (MD), and the mice were then sacrificed. Dissected visual cortices were homogenized, and then astrocytes were sorted by MACS. cDNA libraries were generated from purified RNA of each sample (Ctrl and MD) for sequencing. **(B)** Heatmap-represented gene expression of cell type-specific markers in the whole visual cortex (MACS-) and sorted astrocytes (MACS+, each column represents a single biological replicate). All values were normalized as Reads Per Kilobase of transcript, per Million mapped reads (RPKM). **(C)** Volcano plots showing gene expression changes (blue, decreased, 106 genes; yellow, increased, 84 genes) in the MD group compared to the Ctrl group. **(D-E)** Selected Kyoto Encyclopedia of Genes and Genomes (KEGG) & Gene Ontology (GO) pathways enriched with upregulated (D) or downregulated (E) genes. Dotted lines indicate *P = 0.05. **(F-H)** Quantified z scored log_2_(RPKM) of *Megf10* (f), *Mertk* (g) and *Axl* (h). n=6 from technical duplicates of 3 biological replicates. For (F), *P=0.01572. For (G), *P=0.1935. For (H), P>0.05, n.s., not significant; paired t test. **(I-J)** Representative z-stacked images (I) and quantification of MEGF10 expression in astrocytes (S100b, J) in V1 of the Ctrl and MD groups. **(K-L)** Representative z-stacked images (K) and quantification of MERTK expression in astrocytes (S100b, L) in V1 of the Ctrl and MD groups. **(M-N),** Representative z-stacked images (M) and quantification of AXL expression in astrocytes (S100b, N) in V1 of the Ctrl and MD groups. For (I, K, M), scale bars = 10 µm. n=24 from 4 mice per group. Mean ± s.e.m. For (J), ****P<0.0001. For all comparisons in (L, N), P>0.05, n.s., not significant; Mann‒Whitney t test.

## Astrocytes utilize MEGF10 to eliminate thalamo-cortical synapses during ODP

MEGF10 is a critical receptor for synapse phagocytosis by astrocytes in the developing LGN and adult hippocampus, especially for excitatory synapses (Chung et al., 2013; Lee et al., 2021). To test whether MEGF10-dependent astrocytic phagocytosis is the critical mediator of monocular deprivation-dependent synapse elimination and remodeling, we utilized *Aldh1l1-CreERT2*:*Megf10^fl/fl^* mice to conditionally knock out *Megf10* specifically in astrocytes. We injected tamoxifen into P21 mice for 5 consecutive days to bypass the requirement of MEGF10 in developing retinogeniculate and V1 (Figure 6A). Knocking out astrocyte-specific *Megf10* in V1 was verified by the absence of MEGF10 immunoreactivity in the P35 V1 (Figure 6B).

**Figure 6.**
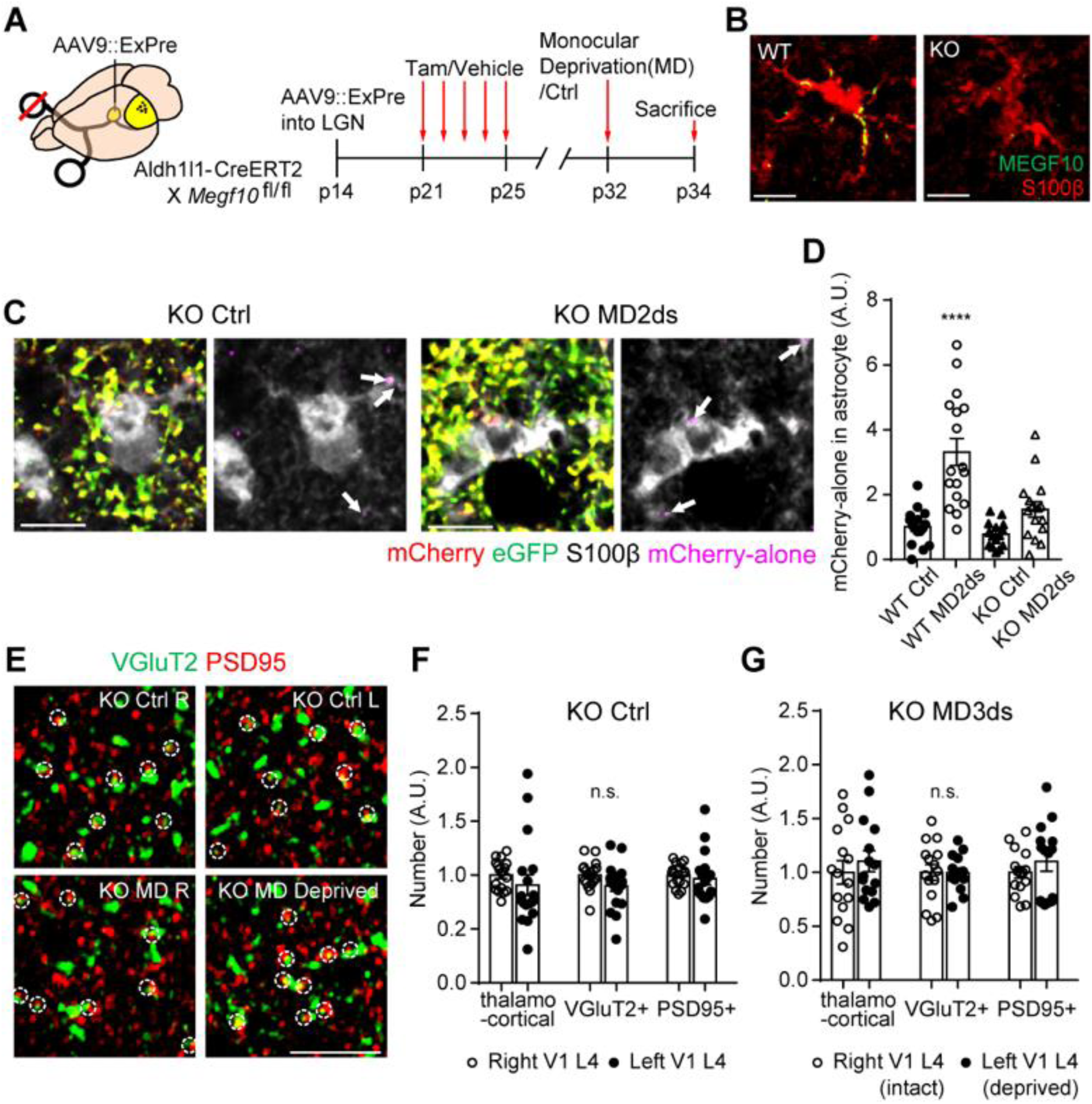
Astrocytes utilize MEGF10 to eliminate thalamo-cortical synapses during ODP. **(A)** Schematic illustration of the experimental design and schedule. All Aldh1l1-CreERT2:*Megf10*^fl/fl^ mice at P14 were injected with AAV9::ExPre into the LGN. Tamoxifen (KO) or vehicle (WT) was injected into the mice for 5 consecutive days from P21. On P32, the right eyes of mice were kept intact (Ctrl) or deprived (MD2ds). All mice were then sacrificed at P34. **(B)** Representative z-stacked images of MEGF10 expression in astrocytes (S100b) in V1 of the WT and KO groups. Scale bars = 10 µm. **(C)** Representative z-stacked images showing ExPre-derived mCherry and eGFP puncta associated with astrocytes (S100b) in V1 of KO Ctrl and KO MD2ds groups. White arrows in the right panels of each group indicate mCherry-only puncta (magenta) inside astrocytes. Scale bars = 10 µm. **(D)** Quantified areas of mCherry-only puncta in astrocytes in V1 normalized to total mCherry-eGFP-labeled synapses. n=15 to 17 from 4 mice per group. Mean ± s.e.m.; (WT MD2ds vs. WT Ctrl, KO Ctrl, and KO MD2ds), ****P<0.0001. Data were compared using two-way ANOVA followed by multiple comparisons. **(E)** Representative single-plane images of thalamo-cortical excitatory synapses (VGluT2, green and PSD95, red) in V1 of the right and left hemispheres of KO Ctrl (KO Ctrl R or L) and MD (KO MD R or Deprived) groups. White dotted circles indicate colocalized synapses. Scale bars = 5 µm. **(F-G)**, Quantified number of thalamo-cortical, VGluT2+ and PSD95+ synapses in V1 of KO Ctrl (F) and KO MD (G) groups. n=15-18 per group from 4 mice. Mean ± s.e.m. For all comparisons in (F, G), P>0.05, n.s., not significant; Mann‒Whitney t test.

To investigate the roles of *Megf10* in the astrocyte-mediated engulfment of thalamo-cortical synapses after monocular deprivation, we injected AAV9::ExPre into the LGN of P14 *Aldh1l1-CreERT2*:*Megf10^fl/fl^* mice and knocked out astrocytic *Megf10* by administrating tamoxifen from P21 to P25 (Figure 6A). Interestingly, when we compared the levels of astrocytic mCherry-only puncta between wild-type (WT) and astrocyte-specific *Megf10* KO mice without monocular deprivation, there was not a large difference between the two groups, indicating that the basal level of astrocytic synapse elimination by MEGF10 in the normal V1 is very low (Figure 6C-6D). However, importantly, we found that the monocular deprivation-induced increase in the level of mCherry-only puncta inside V1 astrocytes was significantly attenuated in astrocyte-specific *Megf10* KO mice (Figure 6D). Although there were no significant differences, the mean value of astrocytic mCherry-only puncta was slightly increased after monocular deprivation even in astrocyte-specific *Megf10* KO mice, suggesting that there could be additional phagocytic receptors in astrocytes other than MEGF10, playing minor roles in eliminating thalamo-cortical synapses (Figure 6D). Taken together, our data show that MEGF10 is a strong driver of monocular deprivation-dependent thalamo-cortical synapse phagocytosis by astrocytes.

Interestingly, when we counted the number of VGluT2- and PSD95-positive excitatory thalamo-cortical synapses after monocular deprivation, we found that astrocyte-specific *Megf10* KO mice after 3 days of monocular deprivation did not exhibit a reduction in the number of VGluT2- and PSD95-positive thalamo-cortical synapses in layer 4 of V1 (Figure 6E-G). These data show that astrocytes utilize MEGF10 to eliminate thalamo-cortical synapses derived from the deprived eye during ODP, and without this astrocytic function, the reduction in synapses in V1 following monocular deprivation fails to occur.

### Astrocyte-mediated synapse elimination is necessary for cortical remodeling

Structural cortical synapse remodeling and plasticity are key features that arise after monocular deprivation (Tagawa et al., 2005). What would be the consequence of failed synapse elimination during ODP? This question has never been adequately addressed since it has not been possible to specifically block the synapse elimination process without affecting general synaptic function and physiology.

Once the contralateral eye is deprived of sensory information, synapses from the intact ipsilateral eye start to dominate visual responses in V1, resulting in the shrinkage of the MZ and the compensatory expansion of the BZ in V1. To compartmentalize the MZ and BZ in V1, we enucleated the contralateral eye 2 hours before sacrificing animals and 3 days after monocular deprivation experiments (Figure 7A). In this experimental setup, BZ V1 neurons that receive inputs from both eyes can be selectively labeled by c-Fos immunoreactivity since MZ V1 neurons that receive inputs from the contralateral eye lose all c-Fos expression due to the complete blockage of the vision-evoked neural activities in the contralateral eye circuit.

**Figure 7.**
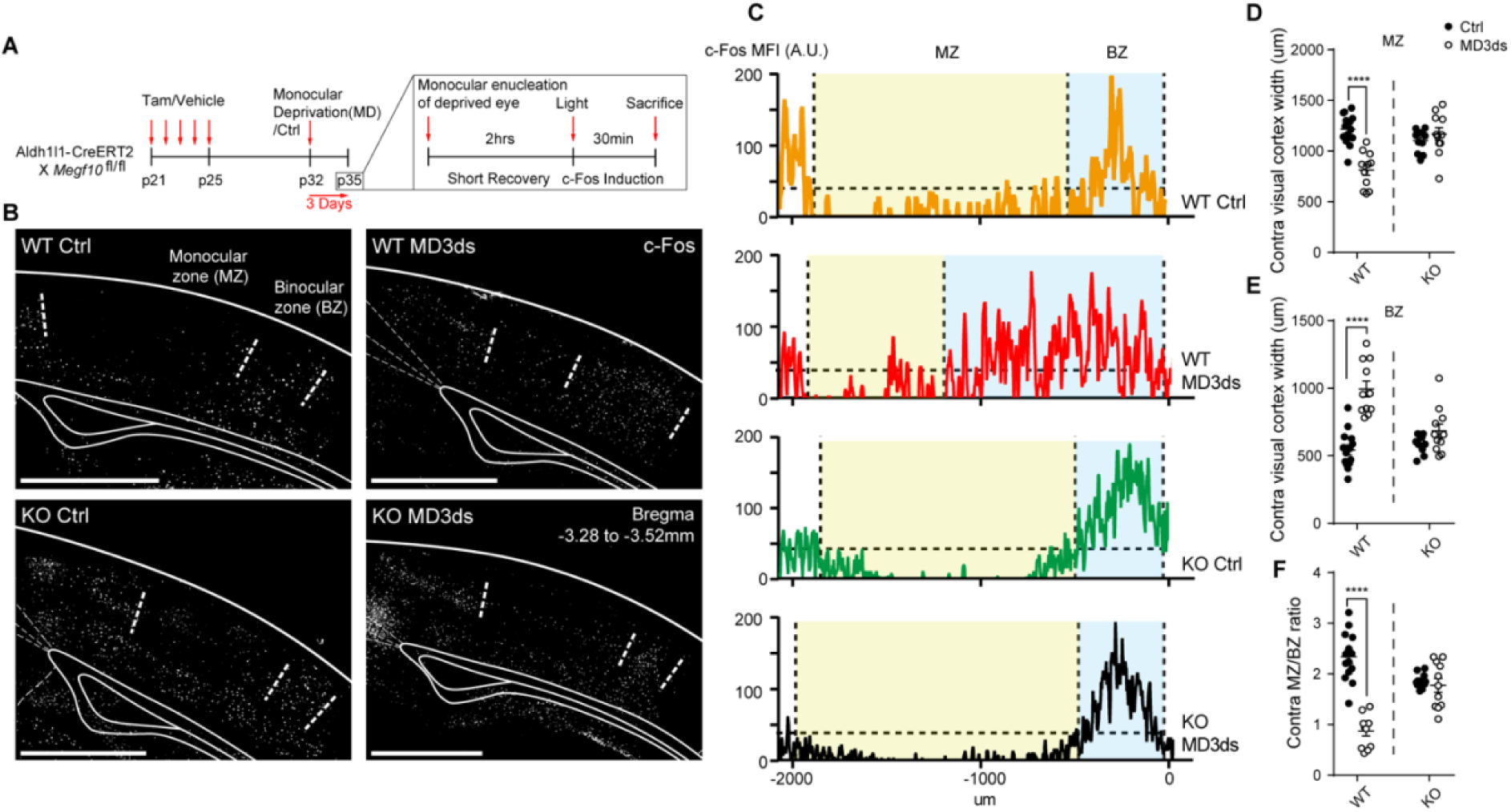
Astrocyte-mediated synapse elimination is required for the expansion of intact synapses. **(A)** Schematic illustration of the experimental schedule. **(B)** Representative z-stacked images of left V1 (Bregma −3.28 to −3.52 mm) with c-Fos staining (white) after enucleating the right eye. White dotted lines compartmentalize the monocular zone (MZ, c-Fos^-^) and binocular zone (BZ, c-Fos^+^). Scale bars = 1 mm. **(C)** Fluorescence trajectories of c-Fos in the V1 of each group. The yellow zone represents MZ, while the blue zone represents BZ. Horizontal dotted lines indicate the threshold of noise. **(D-E)** Quantified widths of MZ (D) and BZ (E). **(F)** Quantified MZ/BZ width ratios. n=11 to 14 from 4 mice per group. Mean ± s.e.m. For (D-F), (Ctrl vs. MD3ds in WT), ****P<0.0001; Mann‒Whitney t test.

In WT mice, we verified that the c-Fos-negative MZ area was shrunken and the c-Fos-positive BZ area was expanded in the contralateral V1 after 3 days of monocular deprivation, indicating that synapses in the MZ and BZ of V1 were successfully remodeled during ODP (Figure 7B-7F). Remarkably, in astrocyte-specific *Megf10* cKO mice, the shrinkage of the MZ as well as the expansion of the BZ were severely compromised, exhibiting almost WT control-like MZ-BZ patterns in V1 even after monocular deprivation (Figure 7B-F). These data strongly indicate that MEGF10-dependent synapse elimination by astrocytes is an essential prerequisite for the subsequent expansion of thalamo-cortical synapses from the nondeprived eye, mediating the completion of the ODP process.

## Discussion

Since Muller and Best’s discovery showing that young kitten astrocytes can restart ODP in the adult cat V1 (Muller and Best, 1989), several recent works have shown that astrocytes participate in ODP-related synapse remodeling by secreting synaptogenic molecules, such as Hevin (Singh et al., 2016) or Chrdl1 (Blanco-Suarez et al., 2018). However, although synapse elimination is another key feature in synapse remodeling, it is unknown how deprived synapses are eliminated in ODP and whether astrocytic phagocytosis participates in this event. By adopting unbiased phagocytosis reporters, we show that astrocytes, but not microglia, start to engulf thalamo-cortical synapses within 24 hours after monocular deprivation. Moreover, through AAV1-mediated transsynaptic tracing, we found that astrocytes preferentially eliminate the deprived synapses, leaving the synaptic connections from the intact eye for subsequent expansion and potentiation procedures. Interestingly, unlike monocular deprivation, binocular deprivation fails to induce astrocyte-mediated synapse elimination, suggesting that the presence of stronger synapses is required for the elimination of weaker synapses, presumably through neuronal/synaptic competition. Finally, we show that deleting astrocytic *Megf10* after P21, which is much later than eye-specific segregation in LGN and developmental synapse pruning in V1, is sufficient to suppress both monocular deprivation-induced synapse elimination and compensatory expansion of synaptic connections from the intact eye.

Although it has been very well established that microglia function as primary phagocytes in developmental synapse pruning, such as in developing the LGN (Stevens et al., 2007; Schafer et al., 2012) and somatosensory barrel cortex (Gunner et al., 2019), recent reports show that the complement and fractalkine pathways, two major pathways for microglia-dependent synapse elimination, are not involved in ODP-related synapse remodeling (Welsh et al., 2020; Schecter et al., 2017). Consistent with these reports, we found that even though microglia appear to possess some portion of mCherry-only puncta in layer 4 of V1, the number of these puncta did not show any changes during ODP, indicating that microglia may not participate in phagocytic elimination of thalamo-cortical synapses after monocular deprivation. Interestingly, it has been shown that microglia exhibit enhanced process motility with an increased synaptic interaction during ODP (Sipe et al., 2016), suggesting potential nonphagocytic roles of microglia in ODP. Recent works have also shown that microglia can directly regulate neuronal activity through adenosine-mediated suppression of the neuronal response (Badimon et al., 2020). Moreover, microglia may clear out the extracellular matrix (ECM) (Nguyen et al., 2020) or interact with astrocytes (Vainchtein et al., 2018; Rothhammer et al., 2018; Liddelow et al., 2017), which may facilitate synapse remodeling processes during ODP. In the future, it will be important to study how astrocytes and microglia interact with each other to achieve precise synapse elimination and subsequent synapse formation during experience-dependent synaptic plasticity.

Intriguingly, our data show that monocular deprivation induces an increase in astrocyte-mediated synapse elimination, while binocular deprivation does not. These results add to the growing evidence that astrocytes recognize and preferentially engulf weak synapses instead of strong synapses, and the presence of strong synapses is required for initiating this elimination process. The exact mechanisms underlying this neuronal/synaptic competition that drives astrocyte-mediated synapse elimination are unclear. However, our previous work in the adult hippocampus also indicates that increasing general hippocampal activity by exposing mice to the environmental enrichment condition (EEC) significantly increases astrocyte-mediated synapse elimination. Thus, our current hypothesis is that the increase in neuronal activity can promote the uncoupling of asynchronous synapses, and that further weakening of synapses during competition may regulate the presentation of synaptic “eat-me” or “don’t eat-me” signals. Previous works from our group and others have shown that phosphatidylserine (PS, Park et al., 2021; Scott-Hewitt et al., 2020) and CD47 (Lehrman et al., 2018) can function as synaptic “eat-me” or “don’t eat-me” signals, respectively, that regulate phagocytic elimination of excitatory and inhibitory synapses via glial cells. Since homosynaptic potentiation can drive heterosynaptic depression in ODP (Jenks et al., 2021), future works will be required to reveal how neuronal activity and resulting homosynaptic/heterosynaptic plasticity regulate the presentation of “eat-me” and “don’t eat-me” signals on a specific population of thalamo-cortical synapses during ODP.

Another interesting result of our study is that monocular deprivation induces the downregulation of astrocytic expression of several key immune components, including complement pathways (*C1qa, b and c*), toll-like receptor pathways (*Ly86*) and chemokine pathways (*CCR5*). These data suggest why microglia do not appear to eliminate thalamo-cortical synapses in V1 during ODP. Although it has been recently shown that astrocytic immune components such as cytokines (e.g., IL-33) are critical modulators of microglial phagocytosis during developmental synapse pruning (Vainchtein et al., 2018), it is still unclear how glial phagocytosis of synapses in turn regulates the CNS immune system. When apoptotic bodies are cleared out by tissue-resident macrophages, macrophages suppress the production of proinflammatory cytokines (e.g., IL-1b, IL-12 and TNF-a) and facilitate the production of anti-inflammatory cytokines (IL-10, TGF-b and PAF, Savill et al., 2002). Since reactive astrocytes express numerous cytokines upon injury and neuroinflammation (Lee et al., 2022), it is plausible that phagocytosis of synapses by astrocytes induces immune tolerance to prevent putative cytotoxic immune responses from occurring in V1 during ODP.

Taken together, our data uncover the novel role of astrocytes in controlling ODP by specifically eliminating deprived thalamo-cortical synapses through MEGF10-dependent phagocytosis, providing answers that have long been sought. Our results also directly show that synapse elimination is an absolutely required prerequisite for the subsequent synapse formation process, such that failures to clear weaker/deprived synapses actively prevent the formation and expansion of strong/intact synapses in ODP. Last, our data suggest that acute modulation of the MEGF10 pathway can be a novel way of manipulating synaptic plasticity in the developing visual cortex. In the future, it would be interesting to study whether such modulation can be useful in preventing ODP and following permanent vision loss due to eye injuries and whether we can also manipulate ODP in the adult brain by astrocytic MEGF10.

## Acknowledgments

We thank all members of the Chung laboratories for helpful discussion. This work is supported by National Research Foundation of Korea (NRF) grants funded by the Korean government (Ministry of Science and ICT, MSIT) 2020M3E5D9079912, 2021R1A2C3005704 and 2022M3E5E8081188 (W.-S.C). J.-H.L was supported by the Global PhD Fellowship Program through the NRF funded by the Ministry of Education (2017H1A2A1042287). Imaris software was provided by the Bio Core facilities of KAIST.

## Author contribution

W.-S.C. and J.-H.L. conceived the project. J.-H.L. designed and performed all key experiments in this study. J.-H.L. analyzed the data. J.K. supported the experiments. J.-H.L. and M.K. performed the bulk RNA-seq experiments. C.-H.S. designed ExPost2 and purified AAV9::ExPost2. W.-S.C. supervised the project. W.-S.C. and J.-H.L. wrote the paper.

## Data availability

A full list of primary antibodies used in this study is in method sections. Recipes for reporters used in this study are in the Methods section. All data and code are available upon reasonable request. For further inquiries, please contact the corresponding author.

---

**Supplementary Figure S1.**
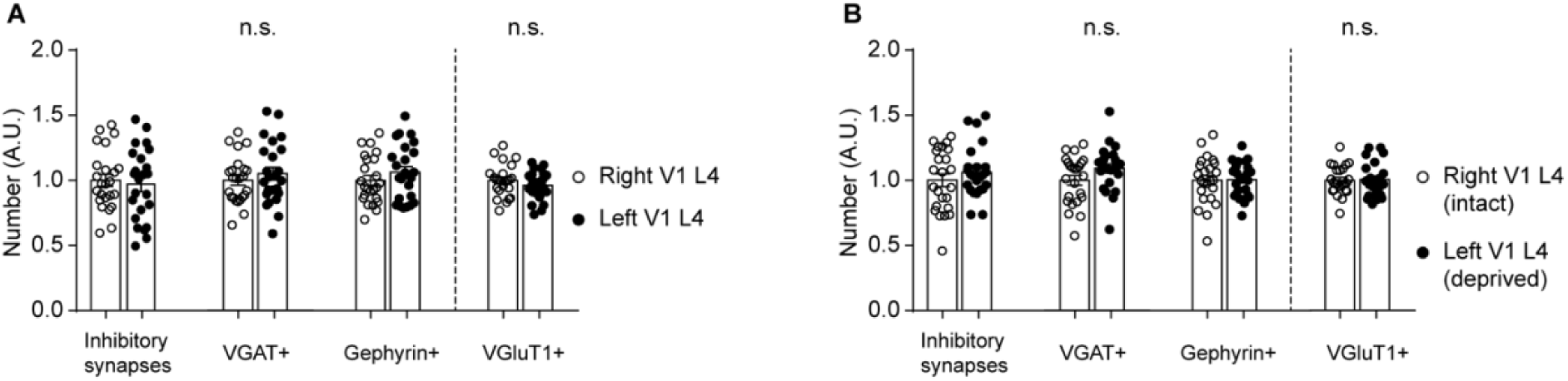
Monocular deprivation does not change the number of inhibitory synapses or VGluT1+ synapses in V1 layer 4. Related to Figure 2. **(A-B)**, Quantified number of inhibitory synapses, VGAT+, Gephyrin+ and VGluT1+ synapses in V1 layer 4 of the Ctrl (A) and MD (B) groups. n=24 per group from 4 mice. Mean ± s.e.m. For all comparisons in (a and b), P>0.05, n.s., not significant; Mann‒Whitney t test.

**Supplementary Figure S2.**
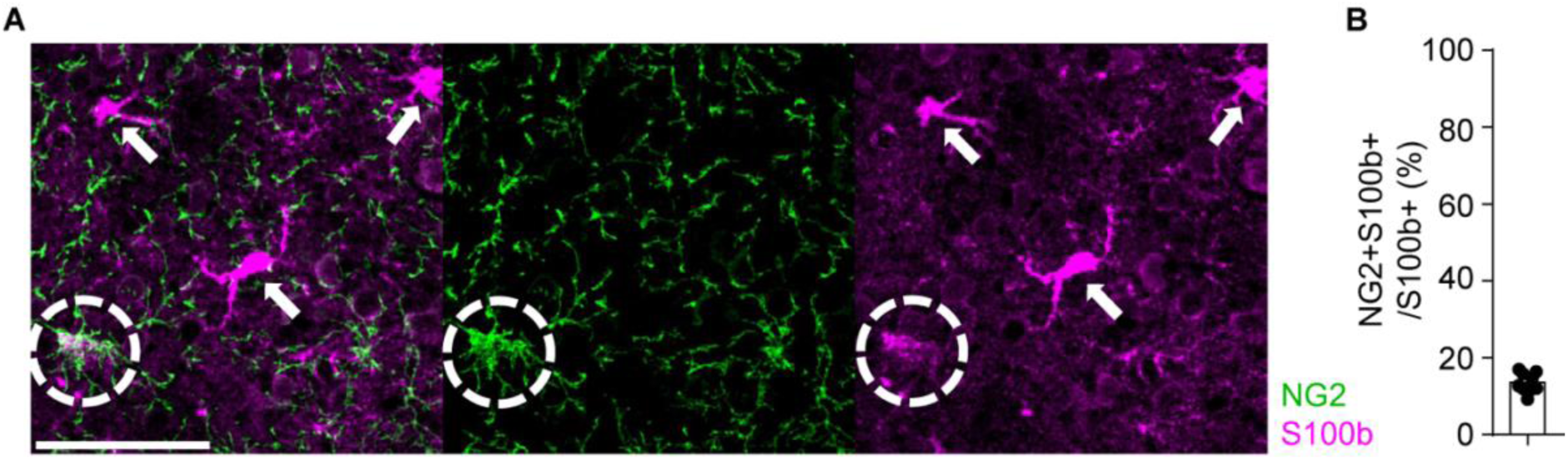
S100b marks the majority of astrocytes with a few oligodendrocyte progenitor cell (OPC) populations. Related to Figure 2. **(A)** Representative z-stacked images of S100b-single positive astrocytes (indicated by arrowheads) and S100b/NG2-double positive OPCs (indicated by dotted circles). Scale bars = 50 µm. **(B)** Quantified percentage of NG2/S100b-double positive OPCs per S100b-positive cells. n=9 from 3 mice. Mean ± s.e.m.

**Supplementary Figure S3.**
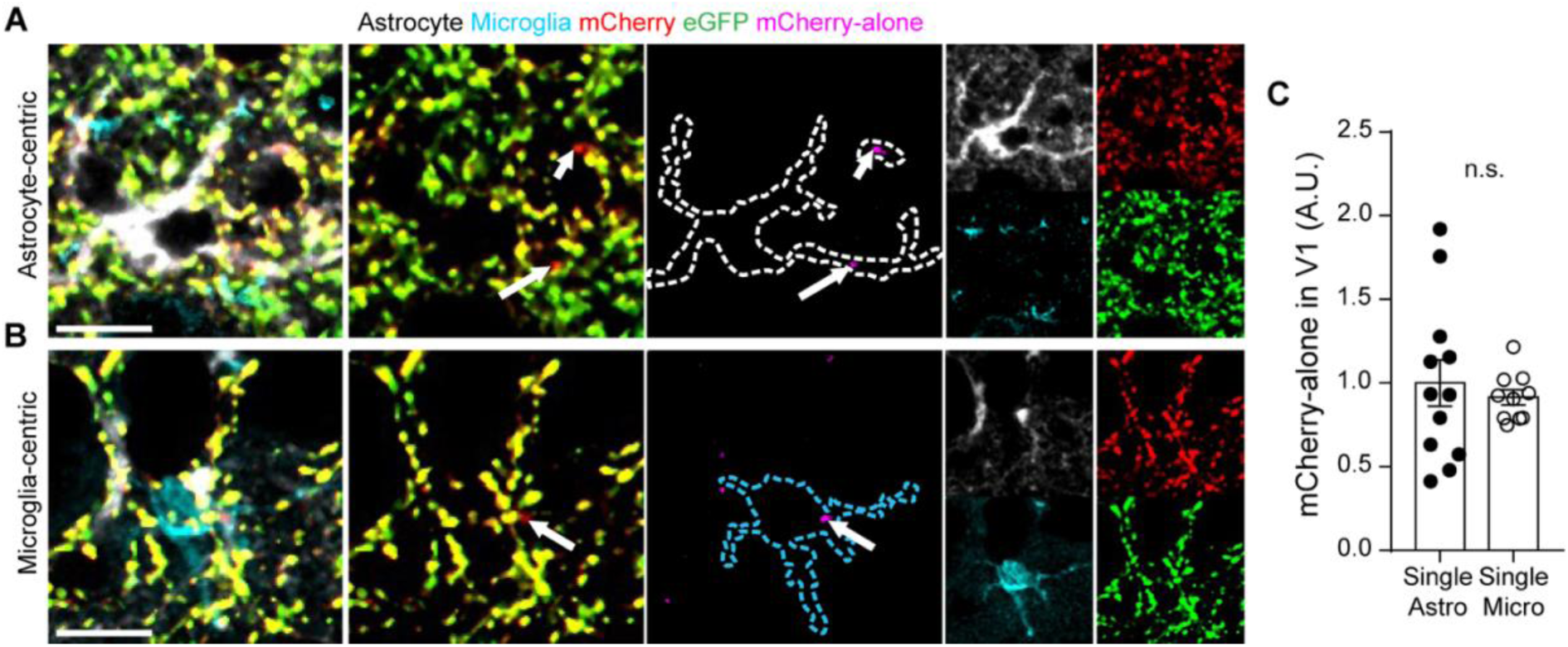
Individual astrocytes and microglia engulf similar levels of synapses in the primary visual cortex. Related to Figure 2. **(A-B)** Representative single plane images of mCherry-eGFP-labeled synapses (green) derived from ExPre in V1 with S100b (white, astrocyte) and IBA1 (cyan, microglia) staining. (A) is astrocyte-centric while (B) is microglia-centric. Scale bar=10 µm. dotted lines highlight the single astrocyte (A, white) or microglia (B, cyan). Arrowheads indicate mCherry-alone puncta (magenta). **(C)** Quantified areas of ExPre-derived mCherry-only in astrocytes and microglia in V1 normalized to total mCherry-eGFP-labeled synapses. n=10 to 12 from 3 mice per group. Mean ± s.e.m. n.s., not significant; Mann‒Whitney t test.

**Supplementary Figure S4.**
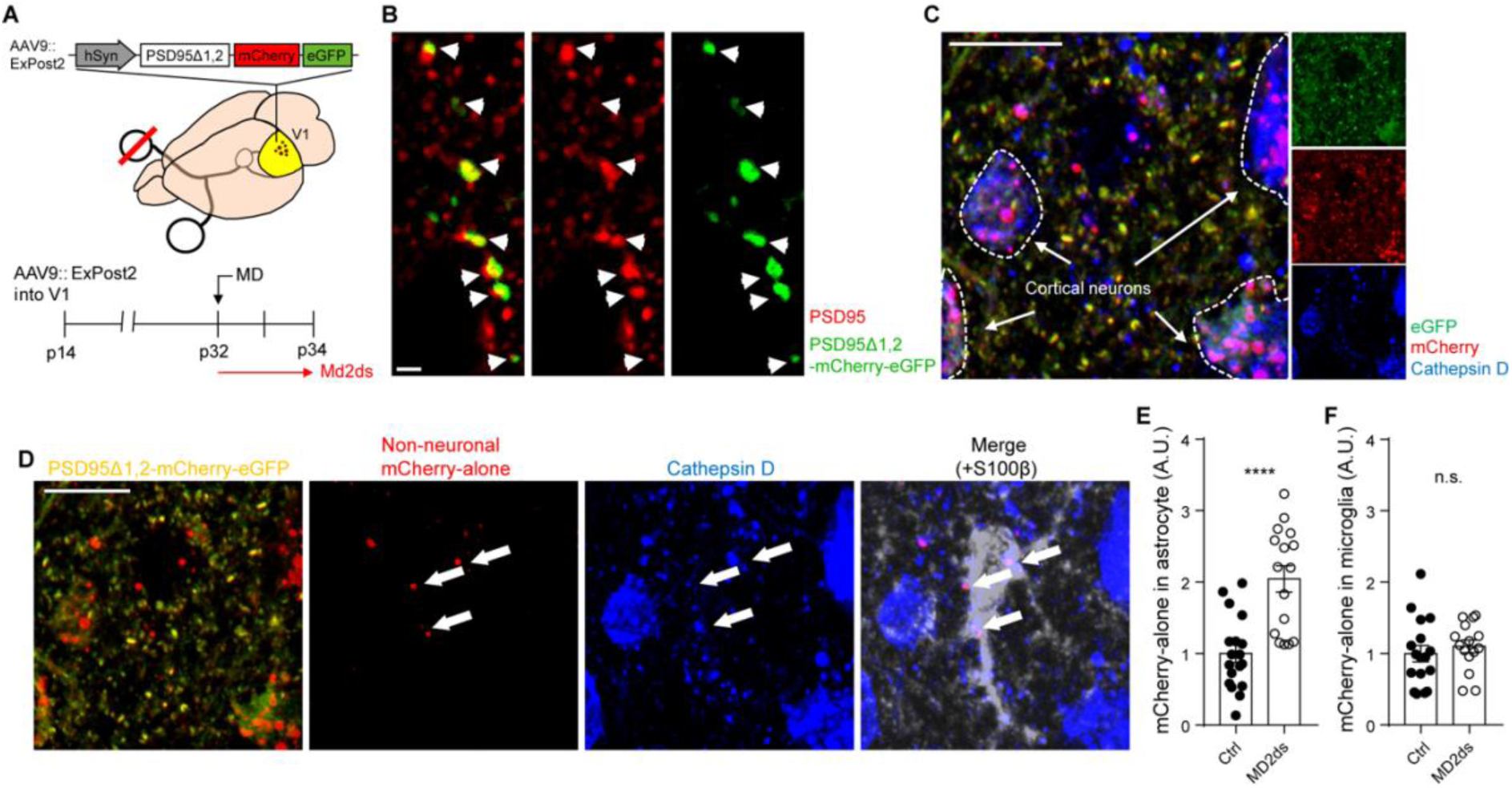
Monocular deprivation induces glial phagocytosis of excitatory postsynapses in V1. Related to Figure 2. **(A)** Schematic illustration of the experimental concept and schedule. Mice at P14 were injected with AAV9::ExPost2 (hSyn-PSDΔ1,2-mCherry-eGFP) into V1. At P32, the right eyes of P32 mice were kept intact (Ctrl) or were deprived, and the mice were sacrificed 2 days later (MD2ds). **(B)** Representative single plane images of mCherry-eGFP-labeled synapses (green) derived from ExPost2 in V1 with PSD95 staining (magenta). White arrows indicate colocalization between mCherry-eGFP and PSD95. Scale bars = 5 µm. **(C)** Representative z-stacked images of mCherry and eGFP expression derived from ExPost2 cells with cathepsin D (blue) staining. Dotted circles indicate cortical neuronal soma. Scale bars = 5 µm. **(D)** mCherry-only puncta derived from ExPost2 (indicated by arrows) are located inside cathepsin D+ astrocytic lysosomes. **(E-F)**, Quantified areas of ExPost2-derived mCherry-only in astrocytes (E) and microglia (F) in V1 normalized to total mCherry-eGFP-labeled synapses. n=11 to 16 from 4 mice per group. Mean ± s.e.m. For (E), ****P<0.0001; for (F), P>0.05, n.s., not significant; Mann‒Whitney t test.

**Supplementary Figure S5.**
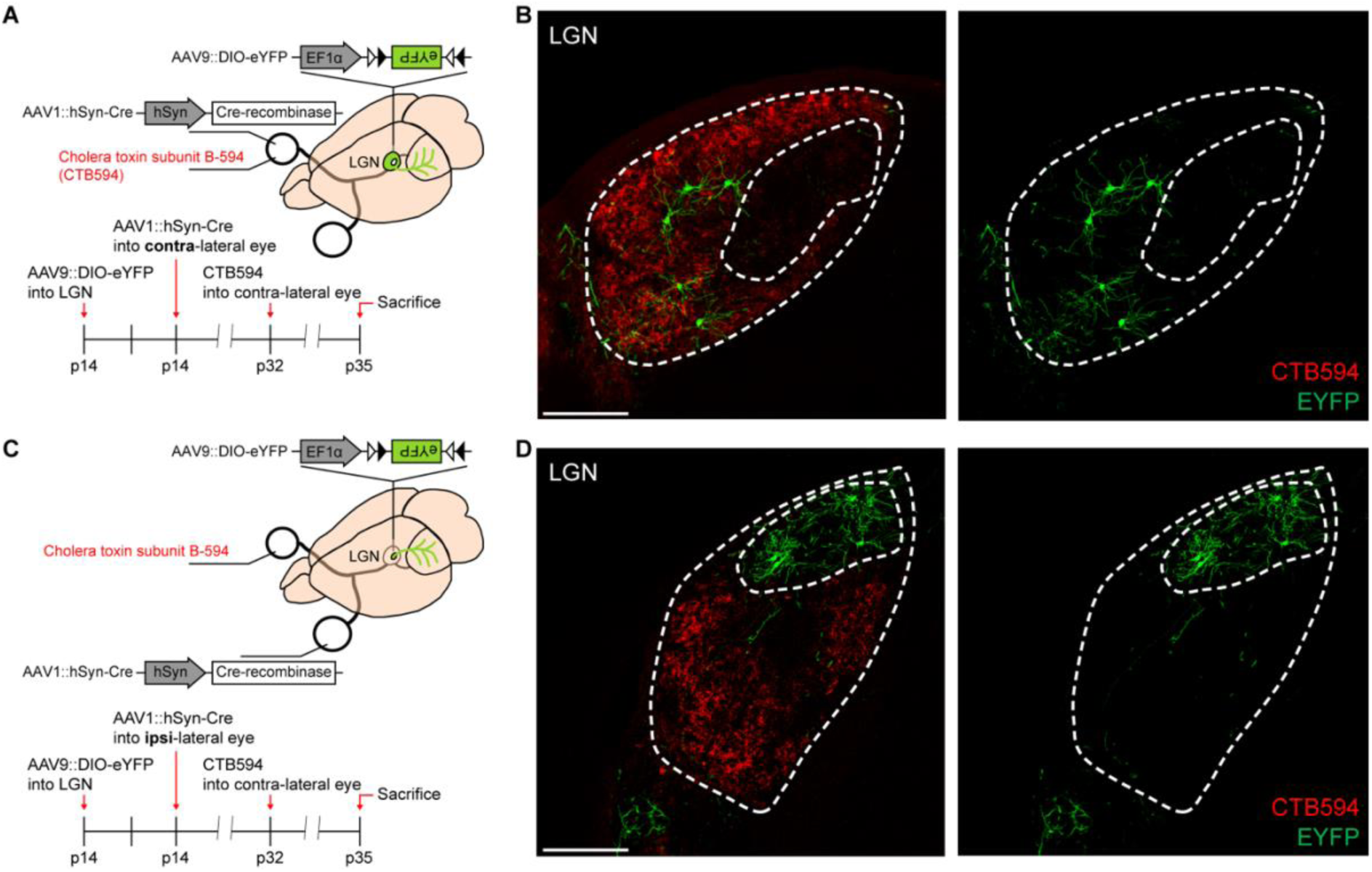
AAV1-mediated transsynaptic tracing enables the labeling of eye-specific thalamic neurons in the LGN. Related to Figure 3. **(A, C)** Schematic illustration of the experimental concept and schedule. At P14, all mice were injected with AAV9::EF1a-DIO-eYFP into the LGN and AAV1::hSyn-Cre into the right (contralateral, A) or left (ipsilateral, C) eye. To label contralateral retinogeniculate synapses in the left LGN, CTB-594 was injected into the right eye at P32. The mice were sacrificed at P35. **(B, D)** Representative z-stacked images of eYFP expression in the left LGN of right eye-injected (B) or left eye-injected (D) mice with AAV1::hSyn-Cre. Scale bars = 5 µm.

**Supplementary Figure S6.**
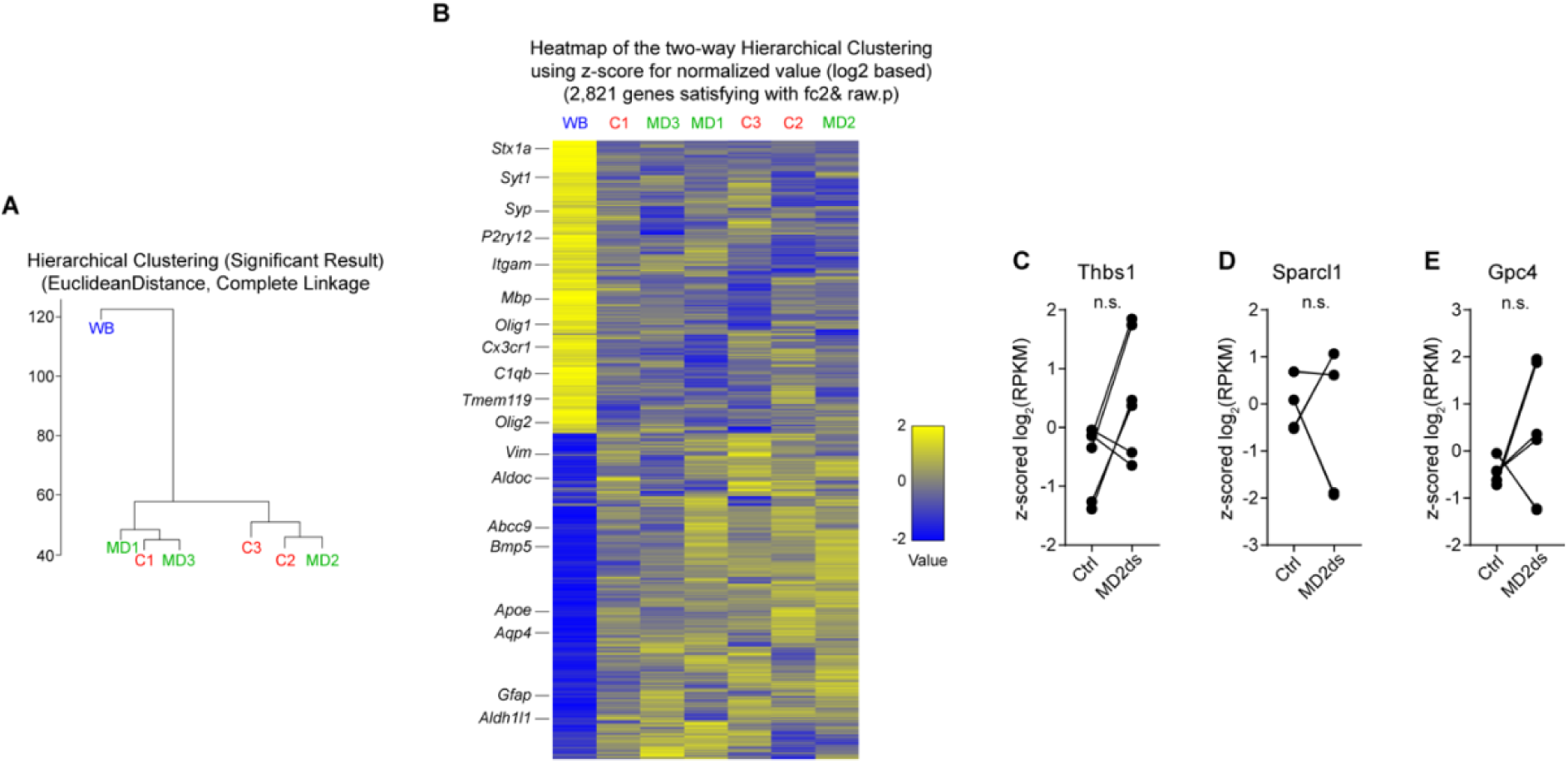
Bioinformatic analysis of astrocytic gene expression in visual cortex after monocular deprivation. Related to Figure 5. **(A)** Hierarchical clustering of gene expression from whole visual cortex homogenate (WB), MACS-enriched astrocytes from control (C1, 2, 3) and deprived groups (MD1, 2, 3) based on Euclidean distance and complete linkage. **(B)** Heatmap of the two-way hierarchical clustering based on z scored values of genes. **(C-E)** Quantified z scored log_2_(RPKM) of *Thbs1* (C), *Sparcl1* (D) and *Gpc4* (E). n=6 from technical duplicates of 3 biological replicates. For all comparisons, P>0.05, n.s., not significant; paired t test.

## Methods

### Mice

All mouse experiments were performed according to articles approved by Institutional Animal Care and Use Committees (IACUC) protocols from Korea Advanced Institute of Science and Technology (KAIST). We followed all proper ethical regulations. LoxP-floxed *Megf10* (*Megf10^fl/fl^*) mice were generated by Stanford Transgenic, Knockout and Tumor Model Center (TKTC). B6;FVB-Tg(*Aldh1l1-Cre/ERT2*)1Khakh/J (*Aldh1l1-CreERT2*) mice were obtained from Jackson Laboratories (JAX). *Megf10^fl/fl^* and *Aldh1l1-CreERT2* lines were crossed to obtain *Aldh1l1-CreERT2*:*Megf10^fl/fl^* (Astrocyte specific Conditional Megf10 Knock-Out) mice. All mouse lines were maintained by breeding with C57BL/6 mice in standard plastic cages (29 cm x 17.5 cm x 18 cm). All experiments involving mutant mice were performed blindly with other littermates. All mice were randomly assigned to experiments. For all experiments, Mus musculus (C57/BL6) was used. Gender was not considered. The mice were sacrificed at the age of postnatal day (P) 32 to P35. For experimental purposes, wild-type C57BL/6 mice were purchased from Daehan BioLink (DBL).

### Plasmids

#### pAAV::hSyn-Cre, pAAV::EF1alpha-DIO-eYFP

pAAV::hSyn-Cre and pAAV::EF1alpha-DIO-eYFP were purchased from Addgene (Addgene items #105553 and #27056).

#### pAAV::hSyn-Synaptophysin-mCherry-eGFP (ExPre)

The plasmid ExPre was generated and utilized according to a previous publication.

#### pAAV::hSyn-PSD**Δ**1,2-mCherry-eGFP (ExPost2)

The truncated version of PSD95 (PSDΔ1,2), which localizes to the excitatory postsynapse with minimal side effects of overexpression (Arnold and Clapham, 1999; Hayashi-Takagi et al., 2015), was synthesized with a gene synthesis service (GenScript). We inserted KpnI and EcoRI restriction sites into the 5’ and 3’ ends of the PSDΔ1,2 sequence by amplifying the PSDΔ1,2 gene with overhang PCR primers containing Kpn1 and EcoR1. The vector (pAAV::hSyn-PSD95-mCherry-EGFP)^27^ was digested with Kpn1 and EcoR1 (Enzynomics) to exclude the PSD95 ORF sequence. The digested PSDΔ1,2 ORF fragment was inserted into the digested pAAV::hSyn-PSD95-mCherry-eGFP (without the PSD95 ORF) by ligation (enzyme manufactured by Enzynomics).

#### pAAV::hSyn-DIO-Synaptophysin-mCherry-eGFP

We inserted NcoI and Asc1 restriction sites into the 5’ and 3’ ends of the Synaptophysin-mCherry-eGFP sequence from pAAV::ExPre by amplification with overhang PCR primers containing NcoI and AscI. After gel extraction, the amplified insert and digested vector (pAAV::EF1alpha-DIO-eYFP) were digested and ligated (Enzynomics, <u>pAAV::EF1alpha-DIO-Synaptophysin-mCherry-eGFP</u>).

We digested pAAV::ExPre and pAAV::EF1alpha-DIO-synaptophysin-mCherry-eGFP with MluI and KpnI. The digested hSyn promoter derived from pAAV::ExPre was inserted into the digested pAAV::EF1alpha-DIO-synaptophysin-mCherry-eGFP (without the EF1alpha promoter) by ligation (Enzynomics).

All cloned plasmids were transformed into Stbl3 (Invitrogen). Candidate plasmids were prepared from inoculated broth by mini prep and sent for validation by a Sanger sequencing service (Bioneer).

### Adeno-associated virus (AAV) production

AAVs were produced following the protocol in our previous publication (Lee et al., 2021). To produce AAVs, we cotransfected a pAAV9 capsid plasmid, a virus assembly helper plasmid (pAd deltaF6, UPENN vector core), and target plasmids into HEK293T (Korean cell line bank) cells using the PEI (1 mg/ml)-based transfection method. pAAV::hSyn-Synaptophysin-mCherry-eGFP (ExPre), pAAV::EF1alpha-DIO-eYFP (DIO-eYFP), and pAAV::hSyn-DIO-Synaptophysin-mCherry-eGFP (DIO-ExPre) were packaged with AAV9. HEK293T cells were maintained with Dulbecco’s modified Eagle’s medium (DMEM, Gibco) containing fetal bovine serum (FBS, Gibco), which was replaced with serum-free media during transfection (6∼18 hours). Transfected cells were incubated in a 37 °C, 5% CO_2_-conditioned cell incubator for 72 hours. Collected cells were resuspended in 50% fresh DMEM and 0.04% DNase I (Worthington) in nuclease-free water and then lysed with a series of freeze‒thaw cycles. AAV-containing supernatant was obtained, and AAVs were further purified by a polyethylene glycol (PEG)-mediated purification method (Lee et al., 2021; Guo et al., 2013). Purified AAVs were concentrated to 200 µl using a 100 kDa Amicon^R^ ultracentrifuge filter tube (Millipore). The titer of AAVs was measured by using AAVpro^R^ titration kit ver.2 (Takara).

### AAV Injection

P14 mice were anesthetized with 1 ml of isoflurane (Piramal) in a sealed plastic box, and their anesthetized state was retained by utilizing a veterinary vaporizer (Surgivet). AAV9::ExPre (5×10^12^ GC/ml) was stereotaxically injected unilaterally into the left LGN (ML: 2.0 mm, AP: −2.1 mm from bregma, DV: −2.55 mm from brain surface) or CA1 (ML: 2.0 mm, AP: −1.9 mm from bregma, DV: −1.4 mm from brain surface). Either AAV9::DIO-ExPre (GC/ml) or AAV9::DIO-eYFP (GC/ml) was stereotaxically injected into the left LGN. AAV9::ExPost2 (6×10^12^ GC/ml) was injected into the left V1 (ML: 2.25 mm, AP: −2.9 mm from bregma, DV: −0.6 mm from brain surface).

All stereotaxic surgeries were performed with a stereotaxic frame (Kopf) and a motorized Hamilton syringe pump (Harvard Apparatus). Prepulled glass pipettes were used as an injection needle (WPI), and glass pipette pulling was performed by a motorized pipette puller (Sutter instrument). After injection, the incision on the head was closed with silk suture (Woori Medical). After surgery, the mice were allowed to recover in a heated cage for an hour before returning to their home cage.

To label eye-specific synapses or axons, anesthetized P14 mice injected with AAV9::DIO-eYFP or AAV::DIO-ExPre into the left LGN were intraorbitally injected with high titered AAV1::hSyn-Cre (2×10^13^ GC/ml, purchased from Vigene Biosciences) into the right or left eye of P14 mice. To label the right eye-derived retinogeniculate axon terminals, CTB-594 (1 mg/ml, Thermo Fisher) was intraorbitally injected into the right eye of the injected mice at P32.

### Monocular/binocular deprivation

To close the eyelid, mice were anesthetized with 1 ml of isoflurane (Piramal) in a sealed plastic box, and their anesthetized state was retained by utilizing a veterinary vaporizer (Surgivet). Mouse heads were restrained in a stereotaxic frame (Kopf). Marginal eyelids were trimmed and sutured with sterile surgical black silk (Ailee Co., Ltd.). During eye closure, the eyeball remained moist by applying sterile saline. After eye closure, ophthalmic ointment (fuscidin) was applied to prevent unwanted infection.

### Immunohistochemistry

Mice were anesthetized with avertin (20 µl/gram) by intraperitoneal injection. Anesthetized mice were perfused with 1x PBS (Welgene) followed by 4% paraformaldehyde in 1x PBS (Wako Chemicals). Brains were isolated and postfixed overnight in fixative at 4 °C. Brains were transferred to 30% sucrose in 1x PBS for 72 hours and then embedded in OCT compound (Leica). Brain sections (40 μm) were prepared by cryo-stat microtomes (Leica). Appropriate sections were collected and embedded in blocking buffer (4% bovine serum albumin, 0.3% Triton X-100 in 1x PBS) for an hour at room temperature (RT). Then, appropriate primary antibodies diluted in blocking buffer were added to brain sections for 24 hours at 4 °C. The details for the primary antibodies used are listed below:

Rabbit anti-S100b (Abcam), rabbit anti-IBA1 (Wako), goat anti-Cathepsin D (R&D systems), guinea pig anti-VGAT (Synaptic systems), guinea pig anti-VGluT2 (Synaptic systems), guinea pig anti-VGluT1 (Millipore), rabbit anti-MEGF10 (Millipore), rabbit anti-PSD95 (Invitrogen), rabbit anti-Gephyrin (synaptic systems), chicken anti-GFP (Aves labs), goat anti-IBA1 (Novus), rat anti-mCherry (Invitrogen), guinea pig anti-S100b (Synaptic systems), rabbit anti-NG2 Chondroitin Sulfate Proteoglycan (Merck), mouse anti-AXL (R&D systems), rat anti-MERTK (eBioscience).

After the brain sections were washed with 0.1% Tween 20 in 1x PBS (PBST), appropriate secondary antibodies conjugated with Alexa Fluor (Invitrogen, Abcam) in PBST were added for 3 hours at room temperature. The sections were washed again with PBST and then mounted on adhesive-coated glass slides (Matsunami or Fisher Scientific). After all sections were fully adhered to glass slides, TrueBlack^R^ (Biotium) diluted to 1/20 in 70% ethanol was added to the sections for two minutes at RT to eliminate autofluorescent signals by lipofuscin. The remaining agent was washed out with PBST, and Vectashield with DAPI or without DAPI (Vector Lab) was used as mounting media. The prepared samples were stored at −20 °C until imaging.

All images were acquired with a Zeiss LSM 880 upright confocal microscope.

### Glial engulfment quantification

Confocal images from the brain sections were acquired by Zeiss LSM 880 (63X or 40X oil immersion optical lens) for quantification as described below. In the graph, each data point represents confocal optical sections containing at least 10∼15 astrocytes. Three to four different data points were acquired from one mouse.

All channels, including the red-colored (mCherry), green-colored (eGFP) and infrared-colored (astrocytes or microglia), were split by ImageJ software. mCherry-only puncta were isolated by subtracting eGFP signals from mCherry signals. Any mCherry puncta overlaid with the minimum eGFP signal were excluded from further quantification and image processing steps. Individual single-plane confocal images from stacks of isolated mCherry-only, green and infrared signals were analyzed. The colocalization assay was performed using the Diana plugin (Gilles et al., 2016).

To prevent complications due to the potentially different expression levels of the AAV synaptic reporters, we converted the images of mCherry-only puncta into black and white binarized images. Then, the area of mCherry-only puncta was measured. This value was further normalized to the area of glial cells (to compensate for potential differences in the density of glial cells) and the area of mCherry-eGFP-labeled synapses (to compensate for potential differences in AAV injection). For Fig. 4, the number of mCherry-only puncta was normalized to the number of nearby mCherry-eGFP-labeled synapses.

### Synapse number quantification

Confocal images of the brain sections were acquired by a Zeiss LSM 880 (63X oil immersion optical lens) for quantification as described below. In the graph, each data point represents confocal optical sections, and 3∼6 different data points were acquired from one mouse.

All channels, including presynaptic and postsynaptic compartments, were split by ImageJ software. A colocalization assay was performed using the Diana plugin described above. The number of presynaptic-only and postsynaptic-only puncta as well as the number of colocalized puncta (both pre- and postsynaptic) were measured by the Diana plugin.

### Tamoxifen formulation

Tamoxifen (Sigma) was dissolved in pure corn oil (Sigma) at a concentration of 20 mg/ml and stored at −20 °C. Before usage, dissolved tamoxifen was heated at 55 °C and then injected intraperitoneally into mice at a dose of 3.75 µl/g. To achieve maximum Cre recombination in target cells, tamoxifen in vehicle was consecutively injected into mice for 5 days (once per day).

### Labeling the monocular/binocular zone

P35 mice were anesthetized with 1 ml of isoflurane (Piramal) in a sealed plastic box, and their anesthetized state was maintained by utilizing a veterinary vaporizer (Surgivet). Mouse heads were restrained in a stereotaxic frame (Kopf). Marginal eyelids were trimmed, and the optic nerve was cut with small scissors. After the eyeball was removed, ophthalmic ointment (fuscidin) was applied to prevent unwanted infection. Then, eyelids were sutured with sterile surgical black silk (Ailee Co., Ltd.). After 2 hours of recovery, the mice were conditioned in a light chamber for 30 minutes and perfused and sampled as described in the Immunohistochemistry section.

Sections containing V1 (Bregma −3.28 to −3.52 mm) were collected and embedded in blocking buffer with 0.5% detergent (4% bovine serum albumin, 0.5% Triton X-100 in 1x PBS) for an hour at room temperature (RT). Then, rabbit anti-c-Fos (Cell Signaling) diluted to 1/200 in blocking buffer was added to brain sections for 72 hours at 4 °C. After the brain sections were washed with PBST (0.1% Tween 20 in 1x PBS), donkey anti-rabbit antibody conjugated with Alexa Fluor 488 (Invitrogen, Abcam) in PBST was added and incubated for 12 hours at 4 °C. The sections were washed again with PBST and then mounted on adhesive-coated glass slides (Matsunami). Vectashield with DAPI (Vector Lab) was used as mounting media. The prepared samples were stored at −20 °C until imaging.

All images were acquired with a Zeiss LSM 880 upright confocal microscope. c-Fos immunoreactivity of z-stacked and stitched whole V1 was converted into fluorescence trajectories through ImageJ.

### Whole RNA sequencing

Mice were sacrificed by decapacitation at P34, 2 days after monocular deprivation. To isolate the primary astrocytes in the visual cortex after monocular deprivation, both visual cortices were excised by sterile surgical scissors. The cortices were mechanically homogenized with a manual glass homogenizer in ice-cold Dulbecco’s phosphate buffered saline (dPBS) and subsequently enzymatically digested with papain for 45 min at 34 °C with 5% CO2 and 95% O2. Microglia and myelin debris were subsequently removed through a mouse anti-CD45 (Invitrogen)-coated dish and myelin removal beads (Myelin Removal Beads II, human, mouse, rat, Miltenyi Biotec), respectively. The homogenate was subsequently mixed with mouse anti-ACSA-2 microbeads, and astrocytes were sorted by magnetic-activated cell sorting MACS (Miltenyi Biotec). RNA was purified by Macrogen using an RNeasy Mini kit (Qiagen) following the manufacturer’s instructions.

The purified RNAs were screened to assess the qualities of total RNA integrity using an Agilent Technologies 2100 Bioanalyzer (or 2200 TapeStation) with an RNA Integrity Number (RIN). Only high-quality RNA with an RIN greater than 7.0 was used for RNA library construction. Qualified RNA were used as templates for generating cDNA libraries using SMARTer Universal Low Input RNA for Sequencing, TruSeq RNA Library Preparation v2 Kit. Qualities of cDNA libraries were screened by using TapeStation D1000 Screen Tape (Agilent).

Whole libraries were then subjected to sequencing with the Illumina platform as specified in the Illumina user guide. The indexed libraries were then subjected to paired-end (2×100 bp) sequencing on the Illumina NovaSeq platform (Illumina, Inc.) by Macrogen Incorporated.

### RNA data analysis

Raw reads were preprocessed to remove low-quality and adapter sequences before analysis, and the processed reads were aligned to the Mus musculus (mm10) genome using HISAT v2.1.047. HISAT utilizes two types of indices for alignment (a global, whole-genome index and tens of thousands of small local indices). These two types of indices are constructed using the same Burrows–Wheeler transform/graph FM (BWT/GFM) index as Bowtie2. Because it uses these efficient data structures and algorithms, HISAT generates spliced alignments several times faster than the widely used programs Bowtie and BWA. The reference genome sequence of Mus musculus (mm10) and annotation data were downloaded from NCBI. Then, transcript assembly of known transcripts was performed with StringTie v1.3.4b48,49. Based on the results, the abundance of transcripts and genes was calculated as read counts per kilobase of exon per million fragments mapped (RPKM) values for each sample. The expression profiles were used to perform additional analyses, such as differentially expressed gene (DEG) analysis. DEGs or differentially abundant transcripts between different groups were filtered through statistical hypothesis testing.

The expression profile determined with the HISAT-StringTie pipeline was used to analyze DEGs between comparable samples. In the data preprocessing step, genes with one or more zero read count values in the samples were excluded. To facilitate comparison between samples without bias, size factors and gene-wise variation were estimated from the read count data, and the read count data were normalized using the relative log expression (RLE) method with the DESeq2 R library. The statistical significance of the differential expression data was determined using the DESeq2 nbinomWaldTest50 with a normalized count. The false discovery rate (FDR) was controlled by adjusting the p value using the Benjamini– Hochberg algorithm. For DEGs with a |fold change|≥2 and raw p <0.05, hierarchical clustering analysis was performed using the complete linkage method and Euclidean distance as a measure of similarity. Gene enrichment analysis was performed based on biologically and functionally known gene sets, such as Gene Ontology, with gProfiler (https://biit.cs.ut.ee/gprofiler/).

For the heatmap, the expression of each gene in log_2_RPKM was applied. The heatmap and line plots representing z scored log2(RPKM) values were made using GraphPad Prism 7.

### Software and statistical analysis

For acquiring confocal images, the Zen (Zeiss) software acquisition system was utilized. For image analysis, ImageJ (NIH) and its plugins (e.g., Diana) were utilized. To reconstruct stacked confocal images into 3D structures, IMARIS (Bitplane) was utilized.

All statistical analyses were performed using GraphPad Prism 7 with 95% confidence. Most imaging data comparing the two groups were analyzed by the Mann‒Whitney test. In the case of imaging data comparing more than three groups, one-way or two-way ANOVA followed by multiple comparisons tests was used. The statistical test used for each experiment is reported in the results. All statistical tests are two-sided.

All experiments were duplicated or triplicated with similar results.

